# A Modular Platform for Purification of Organelle-associated Mitochondria Reveals Functional Specialization at Organelle Contact Sites

**DOI:** 10.64898/2026.08.01.742152

**Authors:** George Maxwell Otto, Akimi Green, Jennifer Cabarrús, Francisco Javier Miralles, Parker Torres, Lorenzo DeLeon, Jane Chea, Daphnee M. Marciniak, Claude Beltejar, Young V. Kwon, Shao-En Ong, David M. Shechner, Yasemin Sancak

## Abstract

Mitochondria perform diverse metabolic and signaling functions, yet how these activities are spatially organized within the mitochondrial network of cells remains poorly understood. Organelle contact sites are spatially restricted hubs that regulate mitochondrial metabolism, signaling, and dynamics, and are therefore well positioned to influence mitochondrial specialization. Investigation of contact site-associated mitochondrial populations has been hindered by a lack of methods to isolate these subpopulations. Here, we develop Organelle Contact-dependent Affinity Purification (ORCA), a workflow for the isolation and analysis of subpopulations of intact mitochondria and associated proteins defined by their organelle contacts. ORCA revealed distinct proteomes for mitochondria associated with the endoplasmic reticulum, lysosomes, peroxisomes, and the Golgi apparatus, demonstrating that organelle contacts define biochemically specialized mitochondrial populations. Focused analysis of Golgi-associated mitochondria showed enrichment of mitochondrial ribosomes and increased mitochondrial translation, revealing an unexpected role for Golgi-mitochondria contacts in regulating mitochondrial protein homeostasis. ORCA also identified the previously uncharacterized Golgi protein KIAA0930/GMO1 as an evolutionarily conserved regulator of oxidative phosphorylation at Golgi-mitochondria contacts. Together, our findings establish ORCA as a broadly applicable approach for investigating the spatial organization of intracellular organelles and reveal organelle contacts as key determinants of mitochondrial specialization.

## INTRODUCTION

Mitochondria perform diverse metabolic and signaling functions, yet how these activities are spatially organized within cells remains poorly understood. Because mitochondria continuously undergo fusion and fission, the mitochondrial network is often viewed as a homogeneous organelle in which proteins and metabolites are rapidly mixed. However, increasing evidence suggests that mitochondrial subpopulations can adopt distinct molecular compositions and functions within the same cell. Spatial specialization has been observed across diverse cellular contexts. ER-associated mitochondria preferentially undergo calcium uptake, mtDNA replication and fission, whereas perinuclear mitochondria exhibit elevated mitochondrial translation, lipid droplet-associated mitochondria show reduced β-oxidation, and neuronal branch-point mitochondria undergo asymmetric division (*1–7*). Functionally and compositionally distinct mitochondrial subpopulations also form in response to changes in cellular bioenergetic demand (*8*). Together, these observations suggest that mitochondrial function is spatially organized within cells. Such organization has been proposed to enable mitochondria to tailor metabolic and signaling activities to local cellular demands while allowing distinct, potentially incompatible functional states to coexist within a single interconnected network. However, how this organization is established and maintained remains unknown.

Mitochondria form membrane contact sites (MCS) with virtually every organelle. MCS are uniquely positioned to generate local mitochondrial specialization because they expose subsets of mitochondria to localized calcium fluxes, lipid transfer, metabolite exchange, and signaling complexes while leaving neighboring mitochondria unaffected. Such communication through MCS plays crucial roles in many aspects of mitochondrial biology, including fusion, fission, and metabolism. Moreover, mitochondria-organelle contacts are dynamic and specific, changing in response to nutrient availability, cell differentiation, and pathologies such as infection, metabolic disease, and cancer (*9–14*).

These observations led us to hypothesize that mitochondria-organelle contacts play an important role in the establishment of specialized mitochondrial subpopulations. Testing this hypothesis has historically been difficult because most existing mitochondria enrichment methods isolate the whole mitochondrial network, and cannot systematically interrogate mitochondria associated with diverse organelles. Consequently, whether different organelle contacts establish specialized mitochondrial subpopulations has remained largely unexplored.

Here, we develop <u>Or</u>ganelle <u>C</u>ontact-dependent <u>A</u>ffinity Purification (ORCA), a workflow that combines contact-dependent biotinylation with mitochondrial purification to isolate and characterize mitochondria associated with an organelle of interest. Applying ORCA to four mitochondrial contact sites revealed distinct contact-specific mitochondrial proteomes, uncovering previously unrecognized biochemical specialization of intracellular mitochondria. In addition, ORCA identified non-mitochondrial proteins selectively enriched at specific organelle contact sites, enabling the discovery of novel regulators of mitochondrial function. Using ORCA, we uncover a role for the Golgi apparatus in regulating mitochondrial translation and identify KIAA0930, which we rename GMO1 (Golgi-associated mitochondrial OXPHOS regulator 1), as an evolutionarily conserved regulator of oxidative phosphorylation at Golgi-mitochondria contacts. Together, these findings establish ORCA as a broadly applicable platform for biochemical and functional characterization of organelle contacts, and identify organelle contacts as regulators of mitochondrial specialization, revealing an important mechanism by which mitochondrial functions are spatially organized within cells.

## RESULTS

### Contact-dependent Biotinylation Enables Purification of Biotin-labeled Mitochondria

We sought to purify mitochondrial populations from different organelle contact sites using the BirA-AviTag system, in which the biotin ligase BirA selectively biotinylates the AviTag substrate in a contact-dependent manner **(Fig. 1A)** (*15–18*). We generated U2OS cell lines that stably co-express AviTag-GFP on the cytosolic surface of mitochondria (AviTag-GFP-OMM) and mCherry-BirA in the cytosol to broadly label mitochondria **(Fig. 1B).** Cells grown in standard media showed robust and constitutive AviTag-GFP biotinylation, a signature which disappeared within two weeks of growth in biotin-depleted media **(Fig. S1A)**. While some cell lines show mitochondrial network fragmentation and failure to proliferate in the absence of biotin, we did not observe gross differences in mitochondrial morphology in U2OS cells **(Fig. S1B)** (*15*). Cell growth rate was not affected, and basal, maximal and ATP-linked mitochondrial oxygen consumption showed only a modest decrease upon prolonged biotin depletion **(Fig. S1C-G)**. Addition of 50 µM biotin to biotin-depleted cells resulted in AviTag biotinylation, which was detectable as early as two minutes after biotin addition **(Fig. S1H).**

**Figure 1:**
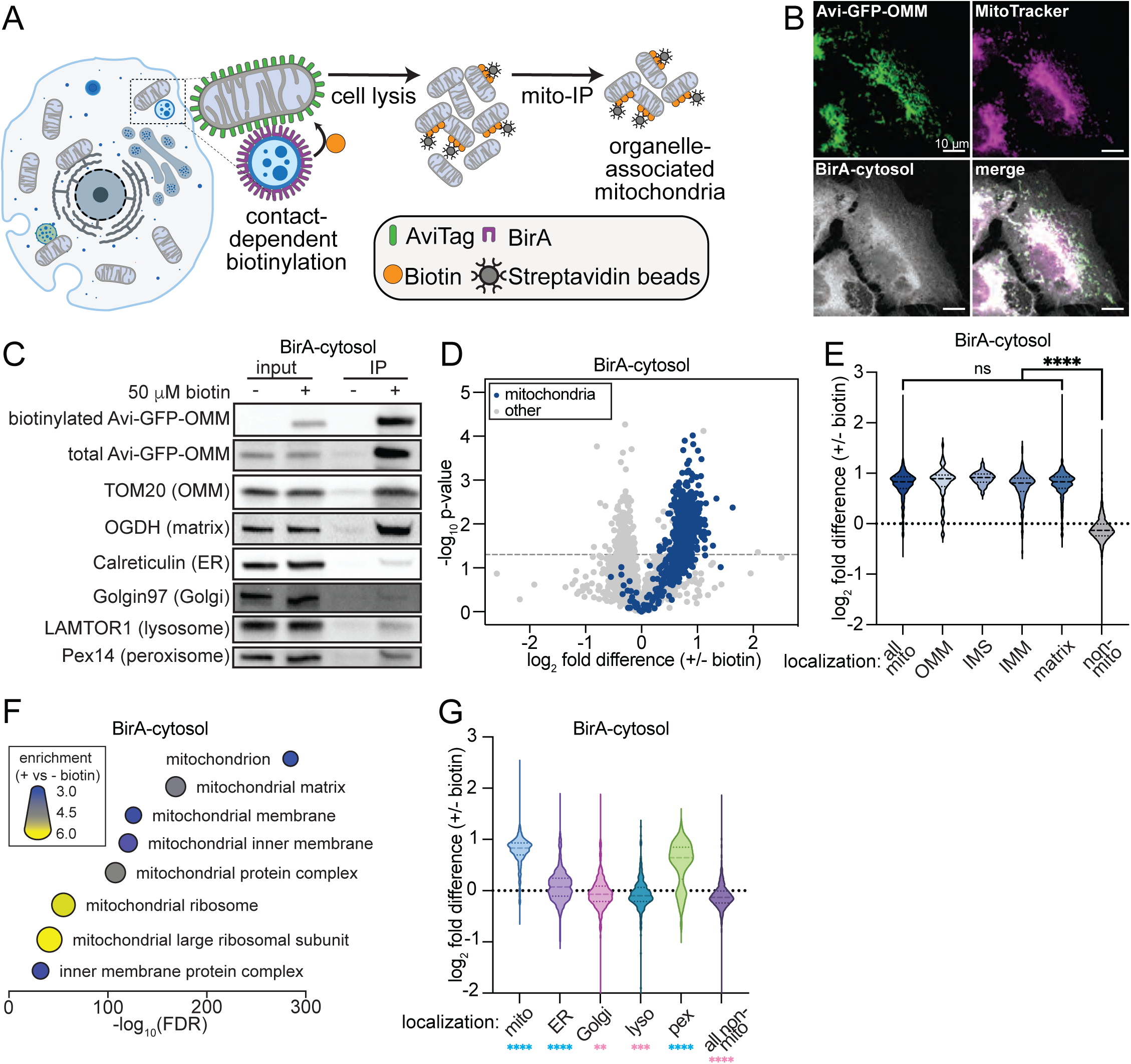
Biotin-dependent labeling and purification of mitochondria. (**A)** Diagram outlining contact-dependent biotinylation of mitochondria-localized AviTag by organelle-localized BirA. **(B)** Confocal microscopy images of MitoTracker Deep Red FM-labeled U2OS cells expressing mitochondrial AviTag-GFP and cytosolic BirA-mCherry. **(C)** Western blot analysis of whole-cell lysates and lysates of streptavidin pulldown taken from the same cells as in **B**, with and without 2 minute incubation with 50 µM biotin. **(D)** Volcano plot showing protein abundance by mass spectrometry from streptavidin pulldown samples prepared as in **C**. Student’s t-test was used to calculate p-values. n=697 mitochondrial and 4651 non-mitochondrial proteins. **(E)** Violin plot showing the distribution of biotin-dependent enrichment values for proteins with the indicated localizations. One-way ANOVA with correction for multiple comparisons was used to calculate p-values. (**F)** Gene set enrichment analysis showing fold-enrichment and FDR for all proteins significantly enriched in +biotin samples compared with all proteins detected. Data generated using the ShinyGO tool (bioinformatics.sdstate.edu). (**G)** Violin plot showing the distribution of biotin-dependent enrichment values for proteins localized to the indicated organelles, with statistics comparing each set to a hypothetical value of 0 by one-sample t-test. Statistical notation in blue indicates biotin-dependent enrichment; notation in pink indicates biotin-dependent depletion. Mito: mitochondria; OMM: outer mitochondrial membrane; IMS: intermembrane space; IMM: inner mitochondrial membrane; ns: p>0.05; *: p<0.05; ****: p<0.0001; FDR: false discovery rate; lyso: lysosome; pex: peroxisome.

After biotin addition, we purified mitochondria from cell homogenates using a modified version of the established mitochondria immunoprecipitation (Mito-IP) protocol (*19*). Western blot analysis confirmed biotin-dependent isolation of intact mitochondria with minimal contamination from other organelles **(Fig. 1C)**. To better characterize the streptavidin Mito-IP samples, we analyzed the samples by mass spectrometry and assessed mitochondrial protein coverage and enrichment. We observed strong coverage (73%) and enrichment (69%) of mitochondrial proteins above background **(Fig. 1D)** (*20*). This remained true regardless of protein sublocalization within mitochondria, indicating that mitochondria remained intact during the purification process **(Fig. 1E)**. Gene Ontology Enrichment Analysis (GOEA) confirmed that mitochondrial proteins were the most strongly overrepresented class in pulldowns after biotin addition **(Fig. 1F)**. Lysosomes, and Golgi proteins were depleted in Mito-IP samples, while ER proteins showed a modest but statistically significant enrichment, consistent with the extensive contacts formed between these organelles, which may be preserved during purification. Among non-mitochondrial organelles, peroxisomal proteins showed substantial enrichment in Mito-IP samples **(Fig. 1G)**. This enrichment was driven by the large fraction of peroxisomal proteins in our dataset (52%; GO:0005777) that also localize to mitochondria and are therefore expected to be strongly enriched in mitochondrial pulldowns **(Fig. S1I)**. Together, these data establish a robust strategy for purification of biotin-labeled mitochondria, providing the foundation for isolating mitochondria in contact with specific organelles.

### <u>Or</u>ganelle <u>C</u>ontact-dependent <u>A</u>ffinity Purification (ORCA) Enriches for Organelle-associated Mitochondria Subpopulations

Next, to better understand how organelle contacts affect the compositional and functional specialization of mitochondria, we targeted mCherry-BirA to the cytosolic surfaces of the ER, peroxisomes, lysosomes and the Golgi apparatus in AviTag-GFP-OMM expressing U2OS cells. Confocal imaging confirmed proper localization of BirA constructs to target organelles **(Fig. 2A)**.

**Figure 2:**
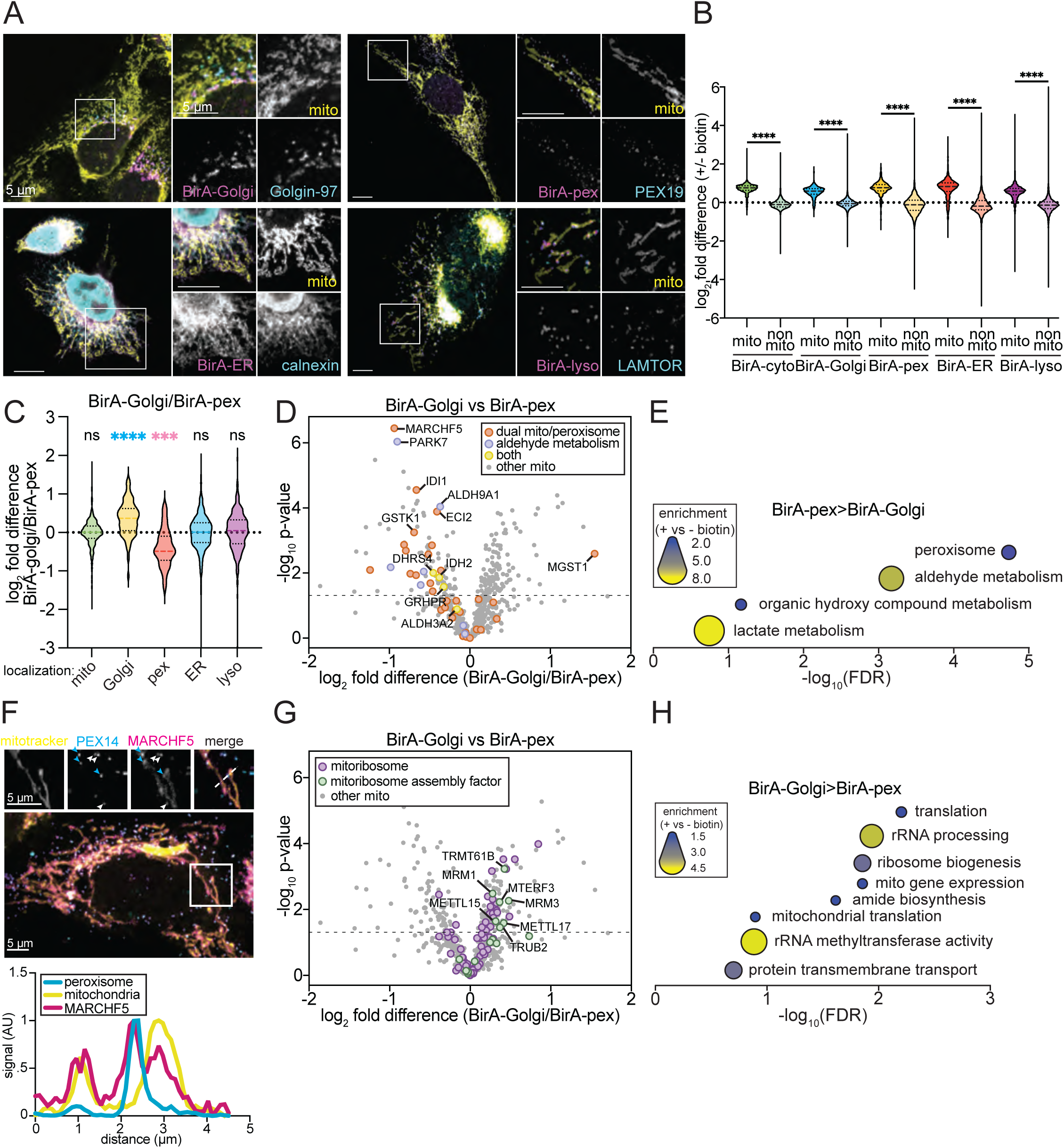
ORCA reveals proteomic differences between mitochondrial subpopulations. **(A)** Confocal microscopy of U2OS cells expressing mitochondrial AviTag-GFP (mito) and the indicated organelle-targeted BirA. Cells were stained with antibodies targeting organelle-specific protein. **(B)** Violin plot showing the distribution of biotin-dependent enrichment values for mitochondrial and non-mitochondrial proteins isolated from U2OS cells expressing mitochondrial AviTag-GFP and the indicated organelle-targeted BirA. Samples were compared using one-way ANOVA with correction for multiple comparisons. (**C)** Violin plot comparing relative protein abundance of organelle proteomes in samples prepared from cells expressing BirA-Golgi or BirA-pex. Samples were compared to a hypothetical value of 0 by one-sample t-test. Statistical notation in blue indicates enrichment in BirA-Golgi samples; notation in pink indicates enrichment in BirA-pex samples. (**D)** Volcano plot showing relative mitochondrial protein abundance in Golgi- and peroxisome-associated mitochondria, with the indicated protein categories enriched in peroxisome-associated mitochondria highlighted. Student’s t-test was used to calculate p-values, n=697 mitochondrial proteins. (**E)** Gene set enrichment analysis showing fold-enrichment and FDR for all mitochondrial proteins significantly enriched in peroxisome-proximal mitochondrial preparations compared with all mitochondrial proteins detected. (**F)** Confocal microscopy and linescan analysis of U2OS cells expressing mScarlet-MARCHF5, labeled with MitoTracker Deep Red FM and stained for peroxisomal protein PEX14. Arrowheads indicate peroxisomes colocalizing with (blue) or distal to (white) mitochondria. **(G)** As in D with the indicated protein categories enriched in Golgi-associated mitochondria highlighted. (**H)** Gene set enrichment analysis showing fold-enrichment and FDR for all mitochondrial proteins significantly enriched in Golgi-proximal mitochondrial preparations compared with all mitochondrial proteins detected. Cyto: cytosol; pex: peroxisome; lyso: lysosome; FDR: false discovery rate. ns: p>0.05; ***: p<0.001; ****: p<0.0001.

We then purified mitochondria from each organelle contact site after biotin addition and calculated the relative abundance of each protein in Mito-IP samples using mass spectrometry. ORCA proteomes were generated in two independent quantitative proteomic series, each including a matched cytosolic BirA control and two organelle-targeted BirA cell lines. Cytosolic controls established mitochondrial enrichment, whereas comparative analyses focused on paired contact-site proteomes acquired within the same experimental series (Golgi versus peroxisome and ER versus lysosome). As with cytosolic BirA, streptavidin pulldown from cells expressing organelle-targeted BirA enriched for mitochondrial proteins **(Fig. 2B)**.

Mitochondria isolated from Golgi- and peroxisome-targeted BirA cells showed modest enrichment of proteins localized to the corresponding BirA-targeted organelle **(Figs. 2C, S2A)**. This indicates that ORCA specifically isolates mitochondria engaged in organelle contacts, and that a subset of these interactions is preserved throughout the purification procedure. This trend was not observed in the ER- and lysosome-associated mitochondrial pulldowns, which instead showed modest but statistically significant enrichment of Golgi, ER and lysosomal proteins in ER-associated mitochondrial preparations. This may reflect the role of the ER in the biogenesis of other organelles in the endomembrane system, or a greater contribution of ER and lysosomal proteomes to the background, resulting in lower signal-to-noise ratio in those preparations **(Fig. S2B)**.

Together, these data establish that ORCA enables purification of organelle-contacting mitochondria and enriches for proteins that interact with mitochondria at the contact site of interest, providing a unique tool and dataset for defining organelle contact-dependent mitochondrial specialization.

### ER- and lysosome-associated Mitochondria Exhibit Distinct Molecular Signatures

ORCA identified 146 mitochondrial proteins with differential abundance between ER- and lysosome-associated mitochondrial populations **(Fig. S2C)**, demonstrating that these abundant mitochondrial subpopulations remain molecularly distinguishable despite extensive overlap within the mitochondrial network. ER-associated mitochondria were enriched in DNA polymerase beta (POLB), which localizes to mitochondria and promotes mitochondrial DNA (mtDNA) repair; mitochondrial rRNA methyltransferase 1 (MRM1), which is positioned near mitochondrial nucleoids; Phospholipid scramblase 3 (PLSCR3), and sterol carrier protein 2 (SCP2), reflecting the role of mitochondria-ER interactions in regulation of mitochondrial DNA synthesis, and lipid homeostasis (*4, 21–27*).

Because ORCA preserves proteins at the mitochondria-organelle interface, we next examined non-mitochondrial proteins recovered with ER- or lysosome-associated mitochondria. The ER proteins VAPB and RRBP1, both implicated as ER-mitochondria tethering factors, were enriched in ER-associated mitochondrial preparations relative to lysosome-associated mitochondria **(Fig. S2D)** (*28, 29*). We also observed enrichment of oxysterol-binding protein 2 (OSBP2), whose yeast homologs mediate transfer of sterols between the ER and mitochondria, suggesting that OSBP2 may perform a similar function in mammalian cells (*30, 31*). Interestingly, lysosome-associated mitochondrial preparations were enriched for glycolysis and gluconeogenesis proteins **(Figs. S2D, E)**. Glycolytic enzymes can assemble into localized metabolons associated with intracellular compartments and mitochondrial outer membrane, and can contribute to the organization of organelle contact sites in plants (*32–34*). This enrichment raises the possibility that mitochondria-lysosome contacts participate in localized metabolic coordination, and glycolytic enzymes may participate in the organization of mitochondria-lysosome contacts.

Together, these findings show that ORCA identifies molecular differences between abundant mitochondrial populations associated with distinct organelle contacts, while identifying proteins residing at the contact interface.

### Peroxisome-associated Mitochondria are Enriched for Proteins Linked to Peroxisomal Metabolism

ORCA identified 200 mitochondrial proteins to be differentially abundant between Golgi- and peroxisome-associated mitochondria **(Figs. 2D, E)**. Peroxisome-associated mitochondrial preparations were enriched for proteins with established dual localization to mitochondria and peroxisomes, including the E3 ubiquitin ligase MARCHF5 and enoyl-CoA delta isomerase 2 (ECI2) (*35–41*). We confirmed localization of MARCHF5 at mitochondria-peroxisome contacts by confocal microscopy **(Fig. 2F)**, supporting the ability of ORCA to identify proteins associated with this contact site.

Mitochondria and peroxisomes are functionally intertwined. They execute complementary steps in the oxidation of very long chain and branched chain fatty acids, necessitating shuttling of metabolic intermediates between the two organelles for full pathway functionality. The two organelles also exchange reactive oxygen species (ROS) through membrane contacts, and both mitochondria and mitochondria-derived vesicles support peroxisome biogenesis (*35, 36, 42, 43*). We analyzed proteins enriched in peroxisome-associated mitochondria to determine if this functional coordination is reflected at the proteome level. GOEA showed enrichment of aldehyde metabolism (GO:0006081), an important component of lipid and amino acid metabolism, in peroxisome-associated mitochondria **(Fig. 2E)**, supporting the idea that peroxisome-associated mitochondria are specialized for metabolic cooperation with peroxisomes.

In addition to mitochondrial proteins, ORCA recovered proteins associated with the peroxisomal membrane, including the biogenesis factors PEX3, PEX13, and PEX16 **(Fig. S2A)**, further demonstrating that proteins residing at the organelle interface remain associated throughout purification, and supporting the role of this contact site in peroxisome biogenesis. Moreover, we identified several proteins with database annotation as both peroxisomal and mitochondrial, but whose putative dual localization has not been experimentally tested **(Figs. 2D-E)**. The broad enrichment of these proteins at mitochondria-peroxisome contacts supports the fidelity of these annotations and is consistent with recent evidence suggesting that dual localized proteins can serve as mitochondria-peroxisome tethers (*44*). Overall, these data show that peroxisome-associated mitochondria are specialized with a unique proteome that supports metabolic crosstalk and organelle biogenesis.

### Golgi-associated Mitochondria are Enriched for the Mitochondrial Gene Expression Machinery

Synthesis of mtDNA-encoded proteins requires coordinated transcription and processing of mitochondrial RNAs and assembly of mitochondrial ribosomes. In mammalian cells, mitochondrial translation produces the 13 mtDNA-encoded subunits of the oxidative phosphorylation complexes (*45*). Surprisingly, Golgi-associated mitochondria showed coordinated enrichment of proteins involved in mitochondrial gene expression, including mitochondrial translation (GO:0032543), rRNA processing (GO:0006364), and mitochondrial ribosome biogenesis (GO:0061668 together with additional assembly factors curated from (*46*)) (**Figs. 2G-H)**. To determine how mitochondrial ribosome abundance in Golgi-associated populations relates to the overall mitochondrial network, we compared Golgi-associated mitochondria with the total mitochondrial pool. This analysis confirmed enrichment of mitochondrial ribosomal proteins in Golgi-associated mitochondria compared to the total mitochondrial pool in the same cell **(Figs. S2F–G)**. We further confirmed this enrichment in an independent cell type. ORCA analysis of HeLa cells similarly identified increased abundance of mitochondrial ribosomal proteins in Golgi-associated mitochondria **(Fig. S2H).**

### Golgi-associated Mitochondria are Characterized by Increased Mitochondrial Translation

The enrichment of mitochondrial ribosomes in Golgi-associated mitochondria suggests that these mitochondria may represent sites of increased mitochondrial protein synthesis. To test this hypothesis, we first examined Golgi-proximal mitochondria by imaging and quantified the abundance of Mitochondrial Ribosomal Protein L9 (MRPL9) and mitochondrial 16S ribosomal RNA (RNR2) in Golgi-proximal (within 1 μm of the Golgi apparatus) and Golgi-distal mitochondria. Both markers were significantly enriched in Golgi-proximal mitochondria **(Figs. 3A-D).** We next measured mitochondrial translation using Fluorescent Noncanonical Amino Acid Tagging (FUNCAT), in which incorporation of the methionine analog HPG into newly synthesized mitochondrial proteins is measured after inhibition of cytosolic translation (*47, 48*). Image analysis showed higher HPG signal in Golgi-proximal mitochondria (within 1 µm of Golgi immunofluorescence signal) after 30 minutes of HPG incorporation **(Figs. 3E-F)**. These findings are consistent with previous reports of elevated mitochondrial translation in perinuclear mitochondria (*1*), and identify Golgi-mitochondria interactions as a likely contributor to this spatial organization.

**Figure 3:**
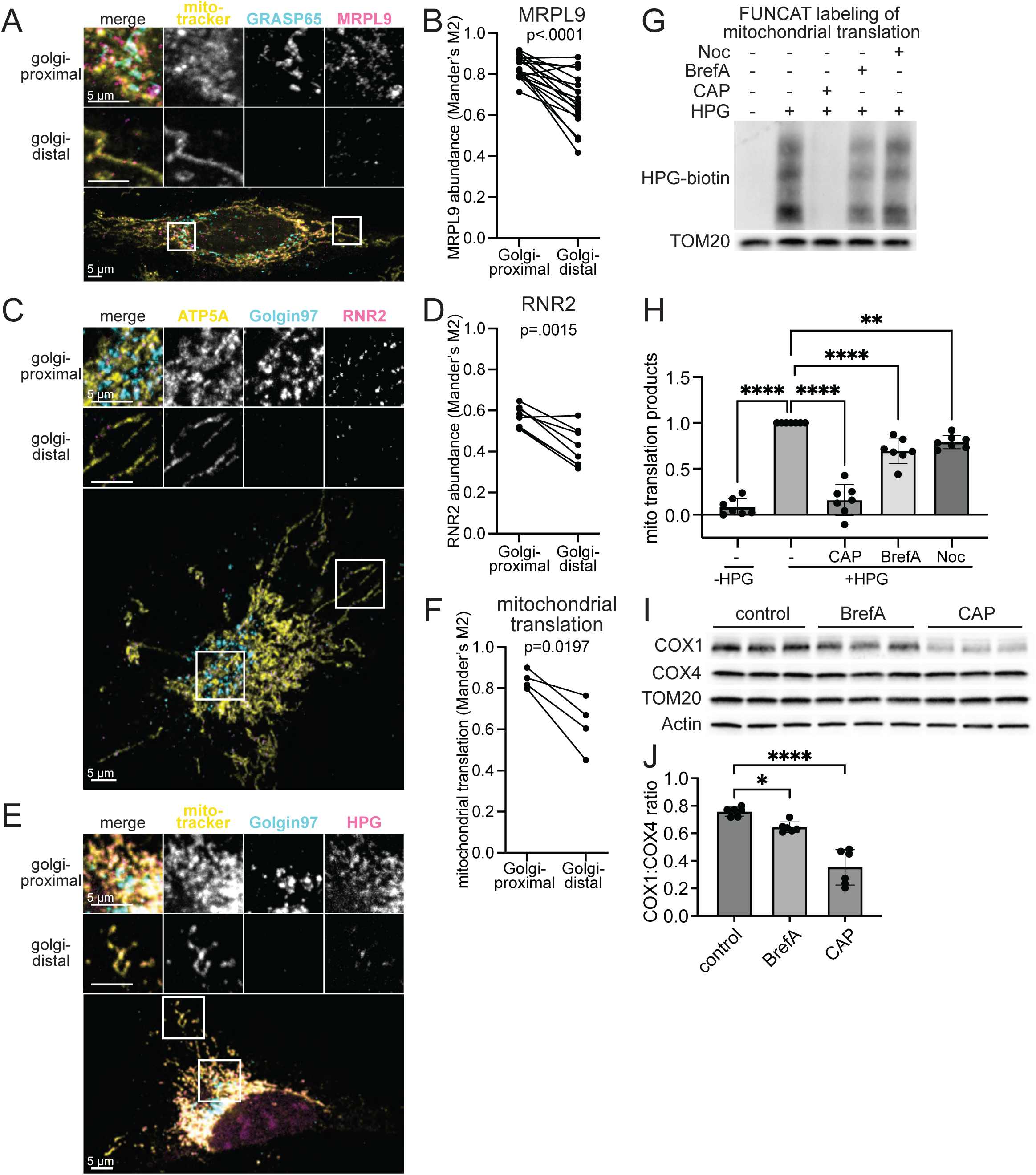
Mitochondrial translation is enriched at Golgi contact sites. **(A)** Confocal microscopy of U2OS cells labeled with MitoTracker Deep Red FM and the indicated antibodies. **(B)** Quantification of images captured as in **A** and compared by Student’s t-test (n=16 cells). **(C)** Confocal microscopy of U2OS cells labeled with the indicated antibodies and RNA FISH oligos targeting RNR2. **(D)** Quantification of images captured as in **C** and compared by Student’s t-test (n=7 cells) **(E)** Confocal microscopy of U2OS cells labeled with HPG and the indicated antibodies. HPG was incorporated into methionine-starved cells in the presence of 100 µg/mL cycloheximide for 30 minutes and conjugated to Picolyl-Azide-AF555 by click chemistry. **(F)** Quantification of images captured as in **E** and compared by Student’s t-test (n=4 cells). **(G)** Western blot analysis of mitochondria-enriched lysates from U2OS cells treated with the indicated drugs. HPG was incorporated into methionine-starved cells in the presence of 100 µg cycloheximide for 2 hours and conjugated to Picolyl-Azide-Biotin by click chemistry. **(H)** Quantification of Western blot samples in **G**. Biotin signal was normalized to TOM20 for each sample and to the matched control sample (+HPG, vehicle) for each replicate (n=7). Statistical comparisons were performed using one-way ANOVA with correction for multiple comparisons. **(I)** Western blot analysis of whole-cell lysate from U2OS cells treated with the indicated drugs for 24 hours. **(J)** Quantification of Western blot samples in **I** (n=6). Statistical comparisons were performed using one-way ANOVA with correction for multiple comparisons. HPG: Homopropargyl glycine, 100 µM; Noc: Nocodazole, 10 µM; BrefA: Brefeldin A, 1 µg/mL; CAP: chloramphenicol, 100 µg/mL. ***: p<0.001; ****: p<0.0001.

To test our hypothesis that Golgi-mitochondria contacts are required to regulate mitochondrial translation, we disrupted their interaction with two different chemicals that interfere with Golgi-mitochondria interactions: Brefeldin A and nocodazole. Brefeldin A blocks vesicular transport from ER to Golgi and leads to collapse of the Golgi apparatus **(Fig. S3A),** whereas nocodazole inhibits microtubule polymerization, resulting in disrupted global organelle distribution and reduced Golgi-mitochondria proximity (*13*). Treatment of cells with either compound for 2 hours reduced mitochondrial translation by ∼30% compared to untreated cells, as measured by biotin-azide labeling of HPG-containing mitochondrial peptides followed by Western blotting. Treating cells with the mitochondrial translational inhibitor chloramphenicol also abolished HPG incorporation, confirming the specificity of this assay to measure mitochondrial translation **(Figs. 3G-H).**

We considered that Brefeldin A treatment could reduce mitochondrial HPG incorporation either through a direct effect on mitochondrial translation or a more general inhibition of global translation. To distinguish between these possibilities, we measured the steady state levels of one mtDNA-encoded and one nuclear-encoded mitochondrial Complex IV subunit (COX1 and COX4, respectively). The COX1:COX4 ratio has previously been used to identify regulators of balanced mitochondrial and nuclear protein synthesis (*49*). Treatment of cells with Brefeldin A or chloramphenicol for 24 hours caused a significant decrease in mtDNA-encoded COX1 protein levels, without affecting nuclear-encoded COX4 levels, suggesting a specific effect of Golgi-mitochondria interactions on mitochondrial translation **(Figs. 3I-J).**

Together, these findings demonstrate that Golgi-associated mitochondria are enriched for mitochondrial translation machinery, exhibit elevated mitochondrial protein synthesis, and require intact Golgi organization to maintain efficient mitochondrial translation.

### Uncharacterized Protein KIAA0930/GMO1 Localizes to the Golgi and Regulates Oxidative Phosphorylation

The finding that Golgi-associated mitochondria exhibit increased mitochondrial translation suggests the presence of Golgi-localized proteins that regulate mitochondrial function. To identify such proteins, we compared our ORCA datasets with a recently published list of empirically defined high-confidence Golgi proteins (*50*). This approach identified KIAA0930, a poorly characterized protein, as the top hit in Golgi-associated mitochondrial pulldowns **(Fig. 4A)**. Based on the findings detailed below, we rename KIAA0930 as Golgi-associated mitochondrial OXPHOS regulator 1 (GMO1), and hereafter refer to it as GMO1.

**Figure 4:**
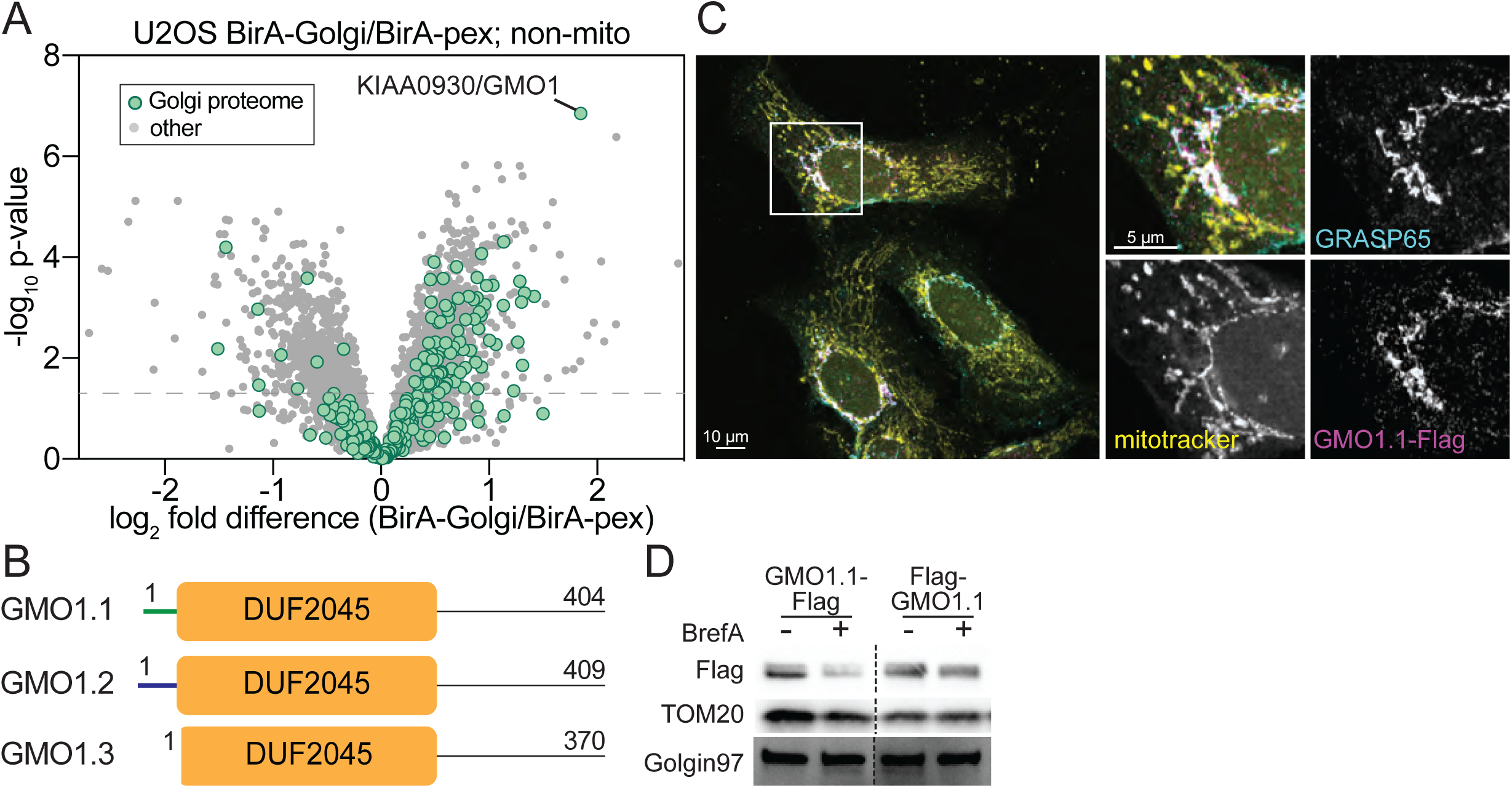
KIAA0930/GMO1 is a novel Golgi protein enriched at mitochondria contact sites. **(A)** Volcano plot showing relative non-mitochondrial protein abundance in Golgi- and peroxisome-associated mitochondria, with putative Golgi proteins defined in Fasimoye et. al. highlighted. Student’s t-test was used to calculate p-values. n=3955 total proteins and 231 putative Golgi proteins. **(B)** Diagram of known GMO1 isoforms with variant regions highlighted in green and blue and the DUF2045 (Domain of Unknown Function 2045) labeled. **(C)** Confocal microscopy of U2OS cells expressing GMO1-Flag (isoform 1) labeled with MitoTracker Deep Red FM and stained with Flag and Golgi protein GRASP65 antibodies. **(D)** Western blot analysis of whole cell lysates from U2OS cells expressing GMO1 isoform 1 with Flag attached to the N- or C-terminus and treated with 1 µg/mL Brefeldin A for 4 hours.

GMO1 is an evolutionarily conserved protein that is defined by a Domain of Unknown Function (DUF2045) **(Figs. 4B, S4A)**. In humans, GMO1 has three isoforms: Isoform 3 consists of 370 amino acids, whereas isoforms 1 and 2 contain an additional exon encoding N-terminal extensions of 34 and 39 amino acids, respectively **(Fig. 4B).** GMO1 is broadly detected across human tissues, with highest expression in the brain and a subset of blood cells (*51*) **(Fig. S4B).** GMO1 was previously identified as a Golgi-localized protein by two independent groups using proteomics of immunopurified Golgi membranes from human cells (*50, 52*) **(Fig. S4C)**; however, its Golgi localization has not been validated by orthogonal means. We tagged all three isoforms with either N- or C-terminal Flag epitopes and expressed them in WT U2OS cells. Isoform 3 (GMO1.3) was expressed at very low levels and was not detected in immunofluorescence experiments, suggesting that it may be unstable under our experimental conditions **(Figs. S4D-E).** Thus, we did not further characterize isoform 3. Both N- and C-terminally Flag-tagged Isoforms 1 and 2 (GMO1.1 and GMO1.2) localized predominantly to the Golgi apparatus **(Figs. 4C, S4D).** Disruption of the Golgi with Brefeldin A treatment markedly reduced abundance of all GMO1 isoforms **(Figs. 4D, S4E),** further supporting a functional link between GMO1 and the Golgi.

Because ORCA captures cytosol-facing proteins associated with mitochondria at organelle contact sites, we reasoned that GMO1 should also face the cytosol. To test this, we expressed cytosolic or Golgi lumen targeted APEX2 in U2OS cells, and labeled proteins in both compartments with biotin (*53*). Biotin-labeled proteins were pulled down and analyzed by Western blotting. Endogenous GMO1 was detected only in pull downs from cytosolic APEX2 expressing cells, indicating that GMO1 is exposed to the cytosol **(Fig. S4F)**. Moreover, GMO1 lacks predicted targeting sequences or high-confidence transmembrane domains, suggesting that it may be peripherally associated with Golgi membranes **(Fig. S4G)** (*54*).

To better understand the function of GMO1, we generated two GMO1 knockout (KO) clones (KO10 and KO20) using CRISPR. GMO1 loss was confirmed by Western blotting **(Fig. 5A).** GMO1 had previously been suggested to localize to mitochondria, based on immunofluorescence data from a proteome-wide screen for protein localization (*55*). However, the reported mitochondrial localization is likely due to nonspecific antibody binding, as we found identical staining between WT and GMO1 KO cells using the same antibody **(Fig. S4H)**. Consistent with this interpretation, Western blotting detected multiple nonspecific bands in addition to GMO1 **(Figs. 5A, S5A)**.

**Figure 5:**
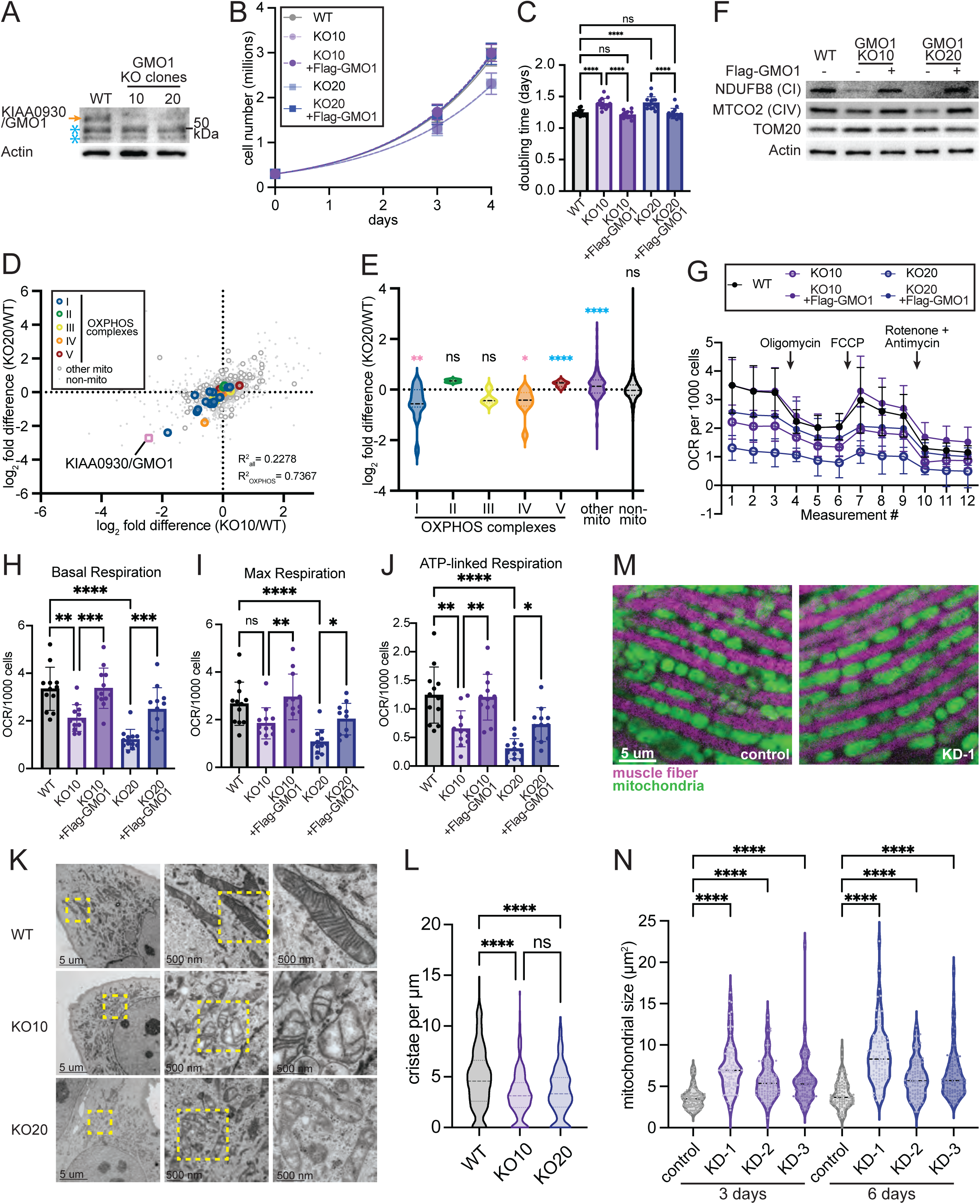
GMO1 is a conserved regulator of mitochondrial metabolism and morphology. **(A)** Western blot analysis of U2OS WT or GMO1 knockout clones. Orange arrow indicates the knockout-validated GMO1 band. Blue asterisk indicates background bands. **(B)** Proliferation assay with U2OS WT and GMO1 knockout clones with and without rescue by re-expression of Flag-tagged GMO1 isoform 2. Statistical comparison was performed using one-way ANOVA with correction for multiple comparisons. n=14 **(C)** Doubling times calculated from proliferation data collected as in **B**. **(D)** Scatter plot showing the abundance for all proteins (n=4639) in U2OS GMO1 knockout clones relative to U2OS WT with select protein categories highlighted. R^2^ was calculated by linear regression in Graphpad Prism. **(E)** Violin plot showing the distribution of fold-change in abundance between GMO1 knockout clone 20 and WT U2OS cells for proteins in the indicated categories. Each sample was compared to a hypothetical value of 0 by one-sample t-test, with pink and blue asterisks indicating significant downregulation or upregulation in GMO1 knockout cells, respectively. **(F)** Western blot analysis of whole-cell lysates from cells of the indicated genotypes. **(G)** Oxygen consumption rate of cells with the indicated genotypes measured by Seahorse assay. **(H)** Basal respiration calculated using the data in **G**. **(I)** Maximal respiration calculated using the data in **G**. **(J)** ATP-linked respiration calculated using the data in **G**. **(K)** Representative electron microscopy images of cells with the indicated genotypes. **(L)** Cristae density quantification for cells imaged as in **K**. **(M)** Confocal microscopy of indirect flight muscle tissue harvested from 3 day-old flies expressing non-targeting control RNAi (*w* RNAi) or RNAi targeting CG9646. **(N)** Quantification of mitochondrial size for samples collected as in **M** with the addition of two additional CG9646 RNAi lines and harvested at 3 and 6 days post-eclosion. CI: Complex I; CIV: Complex IV; OCR: oxygen consumption rate

GMO1 KO clones showed reduced growth compared to WT cells, which was rescued by re-expression of N-terminally tagged GMO1.2 **(Figs. 5B-C).** To define the molecular function of GMO1, we compared the proteomes of WT, GMO1 KO, and rescue cells expressing N-terminally tagged GMO1.2 by quantitative mass spectrometry. GMO1 deletion caused a striking reduction in OXPHOS proteins, particularly Complex I and IV subunits **(Figs. 5D-E, S5B)**. Re-expression of Flag-GMO1.2 restored OXPHOS protein abundance to near WT levels, and rescue cell lines did not show systematic differences in OXPHOS protein expression relative to WT cells **(Figs. S5C-F)**. Notably, enrichment analysis did not identify Golgi-related pathways, indicating that GMO1 loss selectively perturbs mitochondrial proteins despite its Golgi localization. To confirm these observations, we blotted for OXPHOS proteins in GMO1 KO and rescue cell lines expressing untagged GMO1.1, untagged GMO1.2 or Flag-GMO1.2, and detected a consistent decrease in levels of Complex I and IV proteins in KO cells compared to WT and rescue cells **(Figs. 5F, S5G)**. Consistent with these proteomic changes, Seahorse analysis revealed significant reduction in basal, ATP-linked, and maximal oxygen consumption that were restored upon GMO1 re-expression **(Figs. 5G-J)**. Electron microscopy further revealed a dramatic reduction in mitochondrial cristae density in GMO1 KO cells, consistent with impaired assembly or maintenance of the respiratory chain **(Figs. 5K-L)**. Accordingly, fluorescence microscopy showed a significant increase in the number of cells with fragmented mitochondria in GMO1 KO relative to WT and rescue cell lines (**Figs. S5H-I).**

To assess GMO1 function in the context of organism-level physiology and to evaluate its functional conservation across species, we knocked down CG9646, the *Drosophila* ortholog of GMO1, in the indirect flight muscle using three different RNAi lines and imaged mitochondria 3 and 6 days after eclosion. Fluorescence microscopy revealed a dramatic increase in mitochondrial size across all three RNAi lines, a finding that was also supported by electron microscopy imaging **(Figs. 5M-N, S5J-K).** In *Drosophila*, knockdown of NADH:ubiquinone oxidoreductase core subunit S1 (NDUFS1), a core subunit of mitochondrial Complex I, causes mitochondrial enlargement in neurons (*56*). Given that GMO1 knockout reduces the levels of Complex I and IV subunits in human cells, the mitochondrial enlargement induced by CG9646 knockdown may reflect an imbalance in OXPHOS protein levels.

Thus, the development of ORCA enabled identification of KIAA0930/GMO1 as an evolutionarily conserved Golgi protein required for mitochondrial respiratory chain maintenance and mitochondrial ultrastructure. We therefore propose renaming KIAA0930 as Golgi-associated mitochondrial OXPHOS regulator 1 (GMO1).

## DISCUSSION

Despite growing evidence that mitochondrial functions are spatially organized within cells in order to meet local metabolic demands, biochemical characterization of subcellular mitochondria has been limited to tissues where distinct populations can be physically separated (*2, 57–59*). Meanwhile, previous studies of lipid droplet- and endosome-associated mitochondria in human cells reported relatively modest proteomic differences between these populations, leaving open the question of whether organelle contacts generally specify distinct mitochondrial states (*60*). Here, we developed ORCA to biochemically isolate intracellular mitochondrial populations based on their organelle contacts. Applying ORCA to four organelle-mitochondria contact sites revealed different mitochondrial proteomes associated with each contact, providing biochemical evidence that mitochondria are not compositionally uniform but instead comprise a network of spatially distinct functional units, and identifying organelle contacts as a key organizing principle underlying this functional specialization.

Among the contact sites examined, Golgi-associated mitochondria exhibited the most striking specialization, with enrichment of mitochondrial ribosomes and localized mitochondrial translation, revealing a previously unrecognized role for Golgi-mitochondria contacts in mitochondrial protein homeostasis. Interestingly, recent work showed that disruption of Golgi biogenesis proteins GOLGA5 or GOLGA8M alters mitochondrial morphology but has opposing effects on mitochondrial translation, suggesting a complex functional relationship between these organelles (*61*). Our findings provide direct biochemical evidence that mitochondrial translation is spatially linked to Golgi contacts and identify this contact site as one potential regulatory platform.

Identification of GMO1 as an evolutionarily conserved, Golgi-localized regulator of mitochondria highlights the utility of ORCA in discovering mediators of functional and physical organelle interactions. GMO1 homologs are conserved across all major eukaryotic lineages, and are defined by the DUF2045 domain. Previous studies implicated GMO1 and its homologs in cancer in humans, and synaptic vesicle biology in *C. elegans* (*62–67*). However, mitochondrial phenotypes were not previously reported. Our findings identify mitochondrial regulation as a previously unrecognized function of the GMO1 protein family, supported by the altered mitochondrial morphology observed following *Drosophila* knockdown and human knockout experiments. Whether previously reported GMO1-associated phenotypes are due to perturbed mitochondrial function, or reflect divergent GMO1 functions in different tissues and organisms, remains to be seen. In addition, although overexpressed GMO1 exhibited weak cytosolic and nuclear signal in addition to strong Golgi signal, endogenous GMO1 was consistently recovered in Golgi immunoprecipitations, supporting the conclusion that GMO1 is predominantly Golgi localized (*50, 52*). Whether additional pools exist under specific physiological conditions remains an interesting open question for future studies.

Compared with the relatively modest proteomic differences previously reported for lipid droplet- and endosome-associated mitochondria, ORCA identified robust contact-specific proteomes across four intracellular mitochondrial populations (*60*). Several factors could contribute to this difference, including biological variation between contact sites, tissue- or cell type-specific functions, or methodological differences in labeling and purification strategies. Distinguishing between these possibilities will require future comparative studies.

BirA-mediated contact-dependent biotinylation has been successfully used in yeast and human cells (*16, 17*). One limitation of this approach is the need for biotin depletion before labeling, which can disrupt metabolism and cell growth in some cell lines. The recent development of light-activated BirA enables contact-dependent labeling without the need for biotin depletion, and may allow the expansion of ORCA to cell types that cannot tolerate biotin starvation (*15*). More broadly, ORCA provides a modular and generalizable strategy for biochemical interrogation of organelle contacts. Extending this approach to additional organelle subpopulations and physiological contexts will enable systematic dissection of the molecular mechanisms through which organelle interactions spatially organize metabolism and signaling within cells.

## MATERIALS AND METHODS

### Cloning

AviTag-GFP cDNA was synthesized by Twist Biosciences and cloned into the pLYS5 vector (Addgene #50054) between the NheI and EcoRI sites. The OMP25 mitochondrial targeting sequence was appended via Gibson cloning ((*68*); Addgene #83356). BirA constructs were synthesized by Twist Biosciences and inserted into pLVX (Addgene #125839) by restriction cloning. BirA, mCherry, and targeting sequences for organelle-membrane localized BirA constructs are listed in **(Supplemental Table 1)**.

mScarlet (Addgene #189753) and ORFs encoding MARCHF5 (Q9NX47; synthesized by GenScript) or KIAA0930 (isoform 1: Q6ICG6-1, isoform 2: Q6ICG6-2, isoform 3: Q6ICG6-3; synthesized by Twist Biosciences) were combined with the pLYS5 vector by Gibson cloning to make pLYS5-mScarlet-MARCHF5, pLYS5-mScarlet-KIAA0930, and pLYS5-KIAA0930-mScarlet. For KIAA0930 constructs, the mScarlet ORF was further replaced with a Flag epitope via Gibson cloning to generate pLYS5-Flag-KIAA0930 and pLYS5-KIAA0930-Flag.

### Cell Culture and Media

U2OS, HeLa, and HEK293 cells were cultured in DMEM with 4.5 g/L glucose, 1x GlutaMAX (Gibco), and 10% v/v FBS (VWR) or biotin-depleted FBS and grown at 37°C with 5% CO_2_. Biotin-depleted FBS was generated by first incubating 500 mL FBS with 12.5 mL PBS-washed Strep-Tactin beads (IBA) for two hours at room temperature with orbital shaking. Beads were sedimented for 3 minutes at 1000 x g, and the supernatant was subjected to a second two-hour round of shaking with 12.5 mL PBS-washed Strep-Tactin beads. Biotin-depleted FBS was removed from beads, and biotin depletion was confirmed using an IDK Biotin ELISA kit (Immundiagnostik). Biotin concentration was assessed to be lower than the assay’s limit of detection (32.4 ng/L).

### Growth assay

300,000 cells were plated in triplicate in 10 cm dishes and grown in standard conditions for 3 or 4 days, at which point they were collected by trypsinization and counted using a Coulter Z2 Cell and Particle Counter (Beckman Coulter). Doubling time was calculated using the formula *N* = *N*_0_ ∗ 2*^x^*^/*τ*^, where N is the number of cells counted on day x, N_0_ is the number of cells plated on day 0, x is the number of days of proliferation, and *τ* is the doubling time in days.

### Lentivirus Production and Infection

To generate lentivirus, 1×10^6^ HEK293 cells were first plated in a 6 cm dish in standard media and allowed to grow overnight. Viral master mix was prepared by combining 100 ng VSV-G vector (Addgene #8454), 900 ng psPax2 (Addgene #12260), and 1000 ng of the appropriate transgene-encoding lentiviral plasmid in a total volume of 20 µL, adding 150 µL serum-free DMEM, mixing in 6 µL X-tremeGENE transfection reagent (Roche), incubating 15 minutes at room temperature, and adding to HEK293 cells. Two days following transfection, virus was harvested by passing media through a 0.45 µm EZFlow filter (Foxx Life Sciences). Polybrene (8 µg/mL final concentration; Sigma-Aldrich) and 200 µL virus-containing media were added to a 10 cm plate containing the desired cell type at ∼50% confluence. Two days following viral infection, cells were harvested and replated in standard media containing drugs to select for viral integration (1 µg/mL puromycin (VWR) or 100 µg/mL hygromycin B (Sigma-Aldrich)).

KO cell lines were generated using EditCo Human Gene Knockout Kit targeting KIAA0930 with 3 guide RNAs (gRNA) targeting exon 2 (tccaagtccagaagacgatg, ccatttctccatgaagtagg, gatgttctccacctacttca). 40,000 U2OS cells were transfected with 2 µl of Lipofectamine CRISPRMAX Transfection reagent in 25 µl Opti-MEM, 2.6 µl of 3µM gRNA mix and 2 µl of 3µM SpCas9 in 20 µl Opti-MEM supplemented with 2.5 µl of Lipofectamine Cas9 Plus reagent. Cells were mixed with the transfection reagent in duplicate and were plated in 2 wells of a 24-well tissue culture plate. Once confluent, U2OS cells were plated for single cell cloning in 96-well plates in conditioned media (standard media harvested from cells grown for 1-5 days, supplemented with an additional 10% FBS, and filter sterilized). Single cell colonies were established, and genomic DNA was extracted using Qiagen Blood & Cell Culture DNA Mini Kit (#13323). gRNA target region was amplified using a gene-specific primer pair (CACCCTCTTTGATATGCATGAGC; AGCCTCTCTGGATGCTCCA). Gene knockout was confirmed by sequencing.

### ORCA Workflow

Prior to cell harvesting, cells were grown for at least three weeks in DMEM supplemented with biotin-depleted FBS. One million cells were plated in each of five 15-cm dishes and grown to 50-80% confluency. Prior to harvesting, cells were treated with 50 µM biotin or vehicle for two minutes, washed with ice cold PBS, and harvested by cell scraping in 500 µL/plate ice cold KPBS (136 mM KCl, 10 mM KH2PO4, pH 7.25) supplemented with 1x cOmplete mini EDTA-free protease inhibitors (Roche). Cells were pooled and pelleted at 4°C for 1 minute at 1000x g, resuspended in 1 mL KPBS, and lysed on ice with 30 strokes in a 1 mL dounce homogenizer. Homogenates were spun at 4°C for 2 minutes at 1000x g, and supernatants were added to 50 µL KPBS-washed streptavidin beads (Pierce) and incubated on ice for seven minutes with frequent agitation. Beads were collected on a magnet, supernatant removed, and beads were washed 3x with 1 mL KPBS on ice. For Western blot experiments, 50 µL 1x SDS sample buffer (25 mM Tris base, 192 mM glycine, pH 8.3, 0.1% SDS, bromophenol blue) was added to beads and samples were incubated 5 minutes at 95°C and removed from beads before being subjected to SDS PAGE. For proteomics experiments, protein was removed from beads with elution buffer (150 mM NaCl, 50 mM HEPES pH 7.2, 5 mM EDTA, 0.5% SDS, 1x protease inhibitors) for 10 minutes shaking a thermomixer set to room temp and 1500 RPM. Samples were removed from beads and processed as described below.

### Mass Spectrometry

Protein samples were prepared for mass spectrometry analysis following a modified version of the Protein Aggregate Capture (PAC) method (*69*). Solubilized protein samples were aggregated onto KPBS-washed Sera-Mag carboxylate beads (Cytiva) at a bead:protein ratio of 15:1 by adding acetonitrile to the bead:sample mixture at a final concentration of 70% and incubating 10 minutes at room temperature with agitation. Beads were washed with 100% acetonitrile followed by 70% ethanol, resuspended in 100 µL Urea buffer (8M urea, 100 mM Tris 8.5, 10 mM chloroacetamide, 5mM TCEP-HCl), and incubated at 37°C shaking at 1000 rpm for 1 hour. 100 µL TEAB was added to samples along with LysC (NEB) at a ratio of 1:100 LysC:sample protein, and samples were incubated at 37°C shaking at 1000 rpm for 2 hours. 200 µL TEAB was then added to samples along with Trypsin at a ratio of 1:100 Trypsin:sample protein, and samples were incubated at 37°C shaking at 1000 rpm for 16-18 hours. Samples were removed from beads and formic acid was added to a final concentration of 1%. Samples were de-salted on activated C18 stage tips and washed with 50 µL stage tip buffer (5% acetonitrile, 0.1% trifluoroacetic acid).

### Nano-flow liquid chromatography (LC)-MS analysis

Peptide samples were separated on an EASY-nLC 1200 System (Thermo Fisher Scientific) using 20 cm long fused silica capillary columns (100 μm ID, laser pulled in-house with Sutter P-2000, Novato CA) packed with 3 μm 120 Å reversed phase C18 beads (Dr. Maisch, Ammerbuch, DE). The LC gradient was 90 minutes long with 6-45% B at 300 nL/min. LC solvent A was 0.5% (v/v) aq. acetic acid and LC solvent B was 80% acetonitrile in 0.5% (v/v) acetic acid. MS data was collected with a Thermo Scientific Orbitrap Fusion Lumos using a data-independent acquisition (DIA) method with a 120K resolution Orbitrap MS1 scan and 12 m/z isolation window, 30K resolution Orbitrap MS2 scans for precursors from 400-1000 m/z. Data .raw files were converted to .mzML using MSConvert 3.0.21251-d2724a5 and spectral libraries built using MSFragger-DIA (*70*) (w/ FragPipe version 22.0 and MSFragger version 4.1) with quantification through DIA-NN version 1.9 (*71*). The database search was against the UniProt human database (downloaded 2024-01-24) with supplemental spike-in of common contaminants, containing 20477 sequences and 20477 reverse-sequence decoys. For the MSFragger analysis, both precursor and (initial) fragment mass tolerances were set to 20 ppm. Spectrum deisotoping, mass calibration, and parameter optimization were enabled. Enzyme specificity was set to “stricttrypsin” and up to two missed trypsin cleavages were allowed. Oxidation of methionine, acetylation of protein N-termini, −18.0106 Da on N-terminal Glutamic acid, and −17.0265 Da on N-terminal Glutamine and Cysteine were set as variable modifications. Carbamidomethylation of Cysteine was set as a fixed modification. Maximum number of variable modifications per peptide was set to 3. For DIA-NN quantification, FDR was set at 0.01 for global protein group, global precursor, and run-specific precursor FDR. FragPipe/DIA-NN output files were processed and analyzed using the Perseus software package v2.0.11.0 (*72*). Expression columns (protein MS intensities) were log2 transformed and normalized by subtracting the median log2 expression value from each expression value within each MS run. Missing values were imputed from a normal distribution - a width of 2 units and a downshift of 1.8 units was applied. Protein-level differential abundance was assessed using two-sample Student’s t-tests. Benjamini–Hochberg-adjusted false discovery rates (q-values) were also calculated for all quantified proteins.

For MS studies comparing the proteomes of WT, GMO1 KO and rescue cultures; a Quantity.Quality score threshold filter value of .85 was applied to the DIA-NN report.tsv. After empirically confirming all KIAA0930 peptides were excluded from KO samples, precursors were requantified using the MaxLFQ algorithm in the DIA-NN R package 1.0.1. Protein.Group was used as the grouping variable, Precursor.ID as the Identifier, Run as the sample variable, and raw Precursor.Quantity as the input intensity. The curated protein matrix was imported into Perseus, and analyzed as previously described.

### Imaging and Image Analysis

For immunofluorescence imaging, cells were grown to 50% confluency on coverslips (EMS) coated in EmbryoMax 0.1% gelatin (EMD Millipore). Where applicable, mitochondria were labeled with MitoTracker Deep Red (Invitrogen) diluted to 100 nM in OptiMEM (Gibco) for 15 minutes at 37°C 5% CO_2_ prior to fixation. Samples were washed in 1x PBS, and fixed in 4% paraformaldehyde (PFA; EMS) in PBS for 10 minutes. Coverslips were washed thoroughly with 1x PBS and permeabilized with 0.1% Triton X-100 (Sigma) in PBS for 15 minutes rocking on an orbital shaker. Samples were washed again and blocked in 5% BSA in PBS for two hours at room temperature with rocking before incubation with the appropriate primary antibodies diluted in 5% BSA in PBS for 16-18 hours at 4°C with rocking. Samples were washed 4x with 1 mL PBS and incubated with fluorophore-conjugated secondary antibodies diluted in 5% BSA in PBS for 30 minutes at room temperature with shaking. Coverslips were washed 4x with 1 mL PBS and mounted on microscope slides with ProLong Diamond Antifade Mountant (Thermo).

For RNA FISH, samples were processed identically to above until the fixation step. Following fixation, samples were washed twice in glycine quench buffer (0.25 M glycine in 1x PBS) and permeabilized with 0.5% Triton X-100 in PBS for 10 minutes with rocking. Following permeabilization, samples were washed with 1x PBS and incubated in 40% formamide wash buffer (40% formamide, 2x SSC, 0.1% Tween-20) for 5 minutes with rocking. Samples were then incubated with 40% hybridization buffer (40% formamide, 2x SSC, 0.1% Tween-20, 10% dextran sulfate) containing 100 nM primary oligo pool (sequences provided in **Supplemental Table 2**) overnight at 42°C. Samples were washed 3x 5 minutes at 37°C with 30% formamide wash buffer (30% formamide, 2x SSC, 0.1% Tween-20) and incubated with 30% hybridization buffer (30% formamide, 2x SSC, 0.1% Tween-20, 10% dextran sulfate) containing 100 nM secondary oligo (5’–ATGATGATGTATGATGATGTATGATGATGTTTTTTTTT–[AlexaFluor647]–3) for 1 hour at 37°C. Samples were washed 2x 5 minutes in PBST (1x PBS + 0.1% Tween-20) and further processed for immunofluorescence as described above.

Slides were imaged using a Nikon A1R HD25 laser scanning confocal microscope equipped with a Plan Apochromat Lambda 60x oil immersion objective (NA 1.4). Fluorophores were excited using 488, 561, and/or 640 nm lasers as appropriate. For individual experiments, laser power, gain and offset were kept consistent between samples. A pinhole size of 1.2 Airy units, image size of 2048x2048 pixels, z-size of 0.15 µm were used for all images.

Line scan image analysis was carried out in FIJI using the measurement function. To correct for background and facilitate comparison between channels, the minimum value for each channel was subtracted from each value for that channel, and the resulting values were normalized to the maximum value in that channel.

Manders’ M2 coefficient was calculated using Imaris image analysis software. Briefly, Mitochondrial and Golgi structures were defined using the surface tool. Mitochondrial structures were separated into Golgi-proximal (<1 µm from nearest Golgi structure) and Golgi-distal structures. Manders’ M2 coefficient was then calculated to define the proportion of Golgi-proximal and Golgi-distal mitochondria in each cell containing additional signal of interest.

### Cell Lysis and Western Blotting

Cells were grown to 50-80% confluence, washed with ice cold PBS, and harvested in 100-300 µL lysis buffer (150 mM NaCl, 50 mM HEPES pH 7.2, 5 mM EDTA, 1% triton, 1x protease inhibitors) using a cell scraper. Lysate was centrifuged at 4°C 20,000x g for 10 minutes. Protein concentration was determined by Bradford assay (BioRad). Samples were diluted to a final concentration of 1-2 µg/ µL in lysis buffer + 1x SDS sample buffer and incubated at 95°C for five minutes. For experiments in which samples were probed for OXPHOS proteins, this incubation step was instead carried out at 37°C. 8-20 µg sample were loaded onto Tris-Glycine gels (ABclonal) and separated at 120V for one hour before transferring to an Immobilon-PSQ 0.2 µm pore PVDF membrane (Millipore) for 7 minutes at 25V using the Trans-Blot Turbo system (BioRad). Membranes were washed with 1x TBST, blocked with 5% milk in TBST and incubated with the appropriate primary antibodies diluted in 1% BSA in TBST for 16-18 hours at 4°C on an orbital shaker. Following several washes with 1x TBST, membranes were incubated in the appropriate HRP-conjugated secondary antibodies diluted in 1% BSA in TBST for one hour at room temperature. Membranes were washed several times with 1x TBST, incubated with enhanced ECL HRP substrate (ABclonal) and imaged on an iBright CL1000 imager (Invitrogen). Western blot quantification was performed in FIJI using the measure tool with background subtraction. A list of antibodies is provided in **Supplemental Table 3**.

### Metabolic labeling of mitochondrial translation

Metabolic labeling of mitochondrial translation products has been described previously (*47, 48*). Briefly, cells were grown to 50-80% confluency in standard conditions. The morning of the experiment, media was replaced with fresh standard media for one hour, then replaced again with methionine-free media (Gibco Catalog No. 21-013-024 supplemented with 63 mg/L cystine, 10% dialyzed FBS (VWR), and 1x GlutaMax) with 100 µg/mL cycloheximide (Sigma) for 20 minutes. 100 µM homopropargyl glycine (Enzo Life Sciences) was added to the appropriate samples for a further 0.5-2 hours.

For Western blot experiments, cells grown in 10 cm dishes were washed in 1x PBS and harvested in ice cold 500 µL KPBS using a cell scraper and pelleted at 4°C 1000 x g for one minute. Cells were lysed in a dounce homogenizer as described above and clarified at 4°C 1000 x g for 2 minutes. Supernatant was transferred to a new tube and spun at 4°C 10,000 x g for 10 minutes to pellet a mitochondria-enriched membrane fraction. Pellets were resuspended in 60 µL 50 mM Tris 8.8 containing 1x protease inhibitors and 0.4% SDS and incubated at room temperature for 10 minutes with agitation. Lysate was spun at room temperature 10,000x g for 5 minutes and transferred to a new tube where it was mixed with 60 µL click reaction master mix (20 µM Picolyl Azide Biotin (Enzo Life Sciences), 1.2 mM BTTAA, 600 µM CuSO_4_, 5 mM Sodium Ascorbate, 50 mM Tris 8.8, 1x protease inhibitors). Click reaction was allowed to proceed for 1 hour at room temperature. Proteins were then isolated by methanol chloroform precipitation. 480 µL methanol, 120 µL chloroform, and 360 µL MilliQ water were sequentially added to the reaction mixture with vortexing between each step. Samples were spun for one minute at 14,000 x g, the aqueous layer was discarded, and a further 480 µL methanol was added. Samples were vortexed and spun at 20,000 x g for 10 minutes. Liquid was removed from the protein pellet, which was further dried for 20 minutes using a speedvac. Pellets were reconstituted in 100 µL 1x SDS sample buffer for 10 minutes at 37°C and analyzed by Western blot as described above.

For imaging-based experiments, following 30 minutes of HPG incorporation, cells grown on gelatin-coated coverslips in 12-well plates were washed in 500 µL Mitochondria-protective buffer (MPB; 10 mM HEPES, 10 mM NaCl, 5 mM MgCl_2_, 300 mM sucrose, pH 7.5) with 0.005% digitonin, then permeabilized with a further 500 µL MPB + digitonin for 5 minutes. Cells were then washed with 500 µL MPB without digitonin and fixed for 7 minutes in 8% PFA in PBS. Fixed cells were washed 3x with PBS, mitochondrial membranes were permeabilized in 0.5% Triton in PBS, washed again, and blocked with 5% BSA in PBS. Fluorophore was incorporated via click chemistry by incubating with 300 µL click master mix (1.2 mM BTTAA, 1 µM Picolyl-Azide-AF555 (Enzo Life Sciences), 600 µM CuSO_4_, 2 mM sodium ascorbate in PBS) for 40 minutes at room temperature with rocking. Samples were washed 5x with PBS and further processed for immunofluorescence imaging as described above.

### APEX2-mediated biotinylation

2 million U2OS cells that stably express APEX2 in the cytosol (Addgene #215552) or Golgi lumen (Addgene #215554) were plated in 10 cm plates in the presence of 2 µg/ml doxycycline to induce APEX2 expression. Next day, cells were incubated with 5 ml of 500 µM biotin-phenol containing a complete culture media for 30 min at 37°C. Biotinylation was initiated by replacing the media with 4 ml of freshly prepared 1 mM H₂O₂ in PBS for 1 min at room temperature. The reaction was immediately quenched by aspirating the H₂O₂ solution and washing the cells three times with PBS containing 10 mM sodium ascorbate and 10 mM sodium azide. Cells were then washed once with PBS and lysed in 500 µL of 1% Triton X-100 lysis buffer. Biotinylated proteins were isolated using streptavidin-coated magnetic beads. 20 µl of packed beads were used per immunoprecipitation. Beads were washed with lysis buffer prior to use and incubated with clarified cell lysates for approximately 2 h at 4°C with rotation. Following incubation, the beads were washed three times with lysis buffer, and bound proteins were eluted by boiling the beads in 30 µL of SDS sample buffer before SDS-PAGE and immunoblot analysis.

### Conservation analysis

KIAA0930/GMO1 orthologs were identified using OrthoDB and a subset were selected as representatives broadly spanning evolutionary space between humans and algae **(Table 1)**. Representative protein sequences were aligned using Clustal Omega and imported into Jalview to generate residue-level conservation scores.

**Table 1:** GMO1 orthologs.

| **Species** | **Common name** | **NCBI reference seq** | **OrthoDB gene ID** |
| --- | --- | --- | --- |
| S. rosetta | Choanoflagellate | XP_004994312.1 | 946362_0:001276 |
| N. vectensis | Starlet sea anemone | XP_032220552.1 | 45351_0:0021c6 |
| M. musculus | House mouse | NP_001355589.1 | 10090_1:00474f |
| H. sapiens | Human | KAI4003395.1 | 9606_0:002fa6 |
| G. margarita | Arbuscular mycorrhizal fungus | n/a | 4874_0:00144c |
| D. rerio | Zebrafish | NP_001314712.1 | 7955_0:00129a |
| C. arabica | Arabica coffee | XP_071923394.1 | n/a |
| C. elegans | Nematode | NP_509174.1 | 6239_0:004f49 |
| B. bubo | Eurasian eagle-owl | n/a | 30461_0:002321 |
| A. robustus | Anaerobic rumen fungus | ORX77367.1 | 1754192_0:001361 |
| C. primus | Marine green alga | QDZ26062.1 | n/a |

### Electron microscopy

Prior to fixation, U2OS cells were grown to 100% confluency on gelatin-coated ACLAR film (Ted Pella), and *Drosophila* indirect flight muscle was dissected as described below. Samples were fixed with 4% glutaraldehyde in 0.1 M sodium cacodylate buffer and stored at 4°C. Samples were washed 5x for 5 min in 0.1 M sodium cacodylate buffer and post-fixed in buffered 2% osmium tetroxide on ice for 1 hour. Samples were washed 5x in ddH_2_O and subjected to en bloc staining in 1% aqueous uranyl acetate overnight at 4°C. Samples were then washed 5x for 5 min in ddH_2_O, dehydrated sequentially in ice-cold 30, 50, 70, and 95% ethanol and allowed to come to room temperature. This was followed by 2 changes of 100% ethanol and two changes of propylene oxide. Samples were infiltrated in a 1:1 mixture of propylene oxide:Epon Araldite resin for 2 hours followed by 2x 2 hour incubations in fresh Epon Araldite. Samples were then placed in flat embedding molds and polymerized at 60°C overnight. Fixed samples were sliced to a thickness of 80 nm and imaged on a JEOL1230 transmission electron microscope (TEM) at 80 kV.

### Seahorse metabolic flux analysis

Cells were seeded at 30,000 cells per well in a XFe96 Microplate (Agilent Technologies, 103792-100) and cultured overnight in standard conditions. Where applicable, cells were grown in biotin-depleted media for 2-3 weeks prior to experiment start. Extracellular Flux Assay Kit oxygen probes (Agilent Technologies, 103792-100) were activated in XF calibrant buffer (Agilent, 100840-000) at 37°C in a non-CO2 incubator overnight. Seahorse XF medium was prepared fresh by supplementing XF DMEM Medium, pH 7.4 (Agilent, 103575-100) with 1 mM Pyruvate Solution (Agilent, 103578-100), 10 mM Glucose Solution (Agilent, 103577-100), and 2 mM L-Glutamine (Fisher Scientific, BP379-100). Cell culture medium was removed from the microplate, and cells were washed with pre-warmed Seahorse XF medium and refed with fresh Seahorse XF medium. Cells were incubated in a CO2-free incubator at 37°C for 1 hr.

Oxygen consumption rate (OCR) measurements were performed with a Seahorse XFe96 Extracellular Flux Analyzer (Agilent, S7800B) using the Cell Mito Stress Test program. The sensor cartridge was loaded with drugs and inserted into the machine 30 min prior to the flux assay to calibrate oxygen probes. Following three measurements of baseline cellular respiration, respiration was measured three times after addition of each drug: oligomycin, carbonyl cyanide-p-trifluoromethoxyphenylhydrazone (FCCP), and antimycin A + rotenone, with final concentrations of 1 μM for each drug. Oxygen consumption rate was analyzed with the Seahorse XF Cell Mito Stress Test Report Generator (Wave Software).

OCR measurements were normalized to cell number as determined by the CyQUANT Cell Proliferation Assay Kit (Thermo Scientific, C7026). For the CyQUANT assay, media was removed from each well and the plates were frozen at -80°C for at least 1 hr to lyse the cells. CyQUANT was utilized per manufacturer’s instructions to measure fluorescence intensity (Ex 480 nm, Em 520 nm).

### Fly genetics and husbandry

Fly crosses were maintained in vials on standard cornmeal-agar medium and kept at 25°C. We established a recombinant fly line using *UAS-mito-HA-GFP.AP* (BDSC #8442) and *mhc-GAL4.F3-580* (BDSC #38464) to express mitochondrially localized GFP in adult indirect flight muscle. This recombinant fly line was crossed to flies expressing RNAi for CG9646, the *Drosophila* ortholog of KIAA0930/GMO1. For all experiments, newly eclosed male progeny were collected and aged to three or six days. The RNAi lines used in this study: *UAS-w RNAi* (BDSC #28980), *UAS-CG9646 RNAi* (VDRC #103414), *UAS-CG9646 RNAi* (VDRC #14982), and *UAS-CG9646 RNAi* (BDSC #34819).

### Immunofluorescence staining of indirect flight muscle

Mitochondrial GFP was visualized by staining with rabbit anti-GFP polyclonal antibody (1:1000; Proteintech, 50430-2AP) followed by anti-rabbit IgG conjugated to Alexa Fluor 488 (1:2000, Life Technologies, A-11008). Filamentous actin was stained with phalloidin conjugated to Alexa Fluor 594 (1:1000; Thermo Fisher Scientific, A22281). Nuclei were stained with DAPI (1:2000, Sigma-Aldrich, D9542). After aging to three or six days, thoraces of adult male flies were dissected and fixed in 4% paraformaldehyde (PFA) (Electron Microscopy Sciences, P9716) for 20 minutes at room temperature. For permeabilization, we washed samples three times with PBST (PBS supplemented with 0.1% Triton X-100) at 5-minute intervals. For blocking, tissues were incubated in 5% normal goat serum (NGS) (Sigma-Aldrich, G9023) for 1 hour at room temperature. Tissue samples were incubated with primary antibody at the specified dilution overnight at 4°C. The samples were washed three times in PBST at 5-minute intervals. The tissue was incubated with secondary antibody and phalloidin for 1 hour at room temperature in the dark. Samples were then washed three times with PBST at 5-minute intervals, mounted onto microscope slides, and preserved in Vectashield (Vector Laboratories, H-1000). Fluorescence images were acquired using a Leica SP8 laser scanning confocal microscope with 40X/1.25 oil objective lens. Mitochondrial area was measured using ImageJ software.

## Acknowledgements

We thank E. Parker for help with processing samples for EM and image acquisition; Michael Ailion, Amy Clippinger, Suzanne Hoppins and Sancak Lab members for helpful discussions.

## Funding

This work used an EASY-nLC1200 UHPLC and Thermo Fisher Scientific Orbitrap Fusion Lumos Tribrid mass spectrometer purchased with funding from a National Institutes of Health SIG grant S10OD021502 (S-E.O.). This work was supported by Core Grant for Vision Research (NEI P30EY001730), Pew Charitable Trusts (Y.S.), NIH 1DP2ES032761 (Y.S.), Safeway Early Career Award (Y.S.), NIH R21CA288806, R01GM129090 (S-E.O.), NIH R01GM138799-01 (D.M.S.), NIH R01HL160825-01 (D.M.S.), NIH R21HG014903-01 (D.M.S.), NIH R35GM128752 (Y.V.K.),

## Competing interests

The authors declare that they have no competing interests.

## Author Contributions

Conceptualization: Y.S., G.M.O., D.M.S., J.Ca.

Data curation: Y.S., G.M.O., A.G., S-E.O., C.B., D.M.M.

Formal analysis: Y.S., G.M.O., A.G., S-E.O., C.B., D.M.M.

Funding acquisition: Y.S., D.M.S., Y.V.K.

Investigation: Y.S., G.M.O., L.D., A.G., J.Ca., D.M.M., J.Ch., Y.V.K., F.J.M.

Methodology: Y.S., G.M.O., D.M.S., P.T., A.G., S-E.O.

Project administration: Y.S., G.M.O., J.Ca.

Resources: Y.S., D.M.S., S-E.O., J.Ca.

Supervision: Y.S., G.M.O., D.M.S.

Validation: Y.S., G.M.O., P.T., Y.V.K., F.J.M.

Visualization: Y.S., G.M.O., A.G., J.Ca.

Writing – original draft: Y.S., G.M.O., A.G., Y.V.K.

Writing – review & editing: Y.S., G.M.O., A.G., S-E.O., C.B., F.J.M.

---

**Supplemental Figure 1: Biotin-dependent labeling and purification of mitochondria. (A)** Western blot analysis of whole cell-lysates harvested from U2OS cells expressing mitochondrial AviTag-GFP with or without cytosolic BirA-mCherry. Cells were grown in standard or biotin-depleted media for the indicated times. **(B)** Confocal microscopy of U2OS cells expressing AviTag-GFP grown in standard or biotin-depleted media for 3 weeks **(C)** Doubling time calculated for cells grown as in **B** and assessed over a 4-day period. **(D)** Oxygen consumption rate of cells grown as in **B** measured by Seahorse assay (n=24 across 3 biological replicates). **(E)** Basal respiration calculated using the data in **D**. **(F)** Maximal respiration calculated using the data in **D**. **(G)** ATP-linked respiration calculated using the data in **D**. **(H)** Western blot analysis of whole cell-lysate harvested from U2OS cells expressing mitochondrial AviTag-GFP with or without cytosolic BirA-mCherry. Cells were grown in biotin-depleted media for three weeks and treated with 50 µM biotin for the indicated times. **(I)** Violin plot showing the distribution of biotin-dependent enrichment values for proteins localized to the indicated organelles. “Dual loc” indicates proteins with annotated localization to both mitochondria and peroxisomes. Statistical comparisons for all panels are one-way ANOVA with correction for multiple comparisons. ns: p>0.05; *: p<0.05; ***: p<0.001; ****: p<0.0001.

**Supplemental Figure 2: ORCA reveals proteomic differences between mitochondrial subpopulations. (A)** Volcano plot showing relative non-mitochondrial protein abundance in Golgi- and peroxisome-associated mitochondria, with the indicated protein categories enriched in peroxisome-associated mitochondria highlighted. P-values were calculated by student’s t-test. **(B)** Violin plot comparing proteins localized to the indicated organelles for their enrichment in ER-proximal and lysosome-proximal mitochondrial preparations. Samples were compared to a hypothetical value of 0 by one-sample t-test. **(C)** Volcano plot showing relative mitochondrial protein abundance in ER- and lysosome-associated mitochondria, with select proteins highlighted. P-values were calculated by student’s t-test. **(D)** As in C but for non-mitochondrial proteins. **(E)** Diagram of the glycolysis pathway adapted from Cheng et. al. (2013) (*73*) with protein names annotated based on their enrichment and detection. **(F)** Volcano plot showing relative mitochondrial protein abundance in Golgi-associated mitochondria relative to total mitochondrial pool, with select proteins highlighted. P-values were calculated by student’s t-test. **(G)** Violin plot comparing the distribution of enrichment for all mitoribosomal proteins in mitochondrial preparations from U2OS cells expressing BirA-cytosol, BirA-Golgi, and BirA-pex. Statistical analysis shows comparison to a hypothetical value of 0 by one-sample t-test. **(H)** Violin plot comparing the distribution of enrichment for all mitoribosomal proteins in mitochondrial preparations from HeLa cells expressing BirA-cytosol and BirA-Golgi. Samples were compared by one-way ANOVA. Mito: mitochondria; pex: peroxisome; lyso: lysosome; ns: p>0.05; *: p<0.05; ***: p<0.001; ****: p<0.0001.

**Supplemental Figure 3: Mitochondrial translation is enriched at Golgi contact sites. (A)** Confocal microscopy of U2OS cells treated with 1 µg/mL Brefeldin A or an equivalent volume of DMSO for four hours and labeled with MitoTracker Deep Red FM and anti-Golgin-97.

**Supplemental Figure 4: KIAA0930/GMO1 is a novel Golgi protein enriched at mitochondria contact sites. (A)** Conservation analysis of select GMO1 orthologs detailed in Table 1. Top: Conservation score represented as a 10-amino acid rolling average. Bottom: Sequence alignment generated using Clustal Omega and Jalview with each residue color coded according to its conservation score. **(B)** GMO1 transcript abundance across the indicated cell types. Data accessed from the Human Protein Atlas as detailed in Karlsson et. al. (*51*). **(C)** Two-dimensional UMAP generated by Hein et. al. (*52*)showing subcellular compartment clustering based on enrichment scores across a suite of organelle immunocapture experiments. Data accessed from organelles.czbiohub.org. **(D)** Confocal microscopy of U2OS cells expressing the indicated isoforms of GMO1 tagged at the C-terminus (left) or N-terminus (right) labeled with MitoTracker Deep Red FM and stained with Flag and Golgi protein GRASP65 antibodies. **(E)** Western blot analysis of whole cell lysates from U2OS cells expressing the indicated Flag-tagged GMO1 isoforms and treated with 1 µg/mL Brefeldin A for 4 hours. **(F)** Western blot analysis of whole-cell lysates and streptavidin pulldowns from U2OS cells expressing the indicated APEX2 transgene with or without peroxide treatment. **(G)** Predicted probability of membrane topologies for each amino acid in the indicated GMO1 isoform. Data generated using Transmembrane Helix Prediction (TMHMM2.0) accessed at services.healthtech.dtu.dk/services/TMHMM-2.0/ (*54*). **(H)** Confocal microscopy of U2OS WT or GMO1 knockout cells labeled with an anti-GMO1 antibody. nCPM: normalized counts per million; DUF2045: Domain of Unknown Function 2045.

**Supplemental Figure 5: GMO1 is a conserved regulator of mitochondrial metabolism and morphology. (A)** Western blot analysis of U2OS WT or GMO1 knockout clones. Image is a wider crop of the image from **Figure 5A** illustrating abundant background bands. Orange arrow indicates the knockout-validated GMO1 band. Blue asterisk indicates background bands. **(B)** Gene set enrichment analysis showing fold-enrichment and FDR for all proteins enriched in WT U2OS cells compared to GMO1 knockout clones (log_2_FC>0.5 for both WT vs KO10 and WT vs KO20). **(C)** Scatter plot showing the abundance for all proteins (n=4639) in U2OS GMO1 knockout clones relative to the same clones rescued by reintroduction of Flag-GMO1.2 with select protein categories highlighted. **(D)** Violin plot showing the distribution of fold-change in abundance for proteins in the indicated categories between GMO1 knockout clone 20 and the same clone rescued by reintroduction of Flag-GMO1.2. Each sample was compared to a hypothetical value of 0 by one-sample t-test, with pink and blue asterisks indicating significant downregulation or upregulation in GMO1 knockout cells, respectively. **(E)** As in **C** but comparing rescue cell lines to U2OS WT cells. **(F)** As in **D** but comparing GMO1 rescue clone 20 to U2OS WT cells. **(G)** Western blot analysis of whole-cell lysates from GMO1 knockout cells with and without re-expression of the indicated GMO1 isoform. **(H)** Confocal microscopy with cells of the indicated genotypes expressing mitochondrially targeted mCherry**. (I)** Categorization of mitochondria morphologies for cells imaged as in **H.** Conditions are compared via Chi-squared analysis. **(J)** Electron microscopy images of indirect flight muscle from flies expressing either CG9646-targeting RNAi or a non-targeting (*w* RNAi) control with mitochondria pseudocolored in green**. (K)** Quantification of mitochondrial area from images captured as in **J.** Conditions are compared by one-way ANOVA. ns: not significant (p>0.05); *: p<0.05; **: p<0.01; ****: p<0.0001. 1UT: GMO1.1 untagged; 2: 2UT: GMO1.2 untagged; 2N: Flag-GMO1.2

## Notes

### Competing Interest Statement

The authors have declared no competing interest.

