## Supplemental Figures and Tables for "A Modular Platform for Purification of Organelle-associated Mitochondria Reveals Functional Specialization at Organelle Contact Sites"

### Supplemental Figure 1

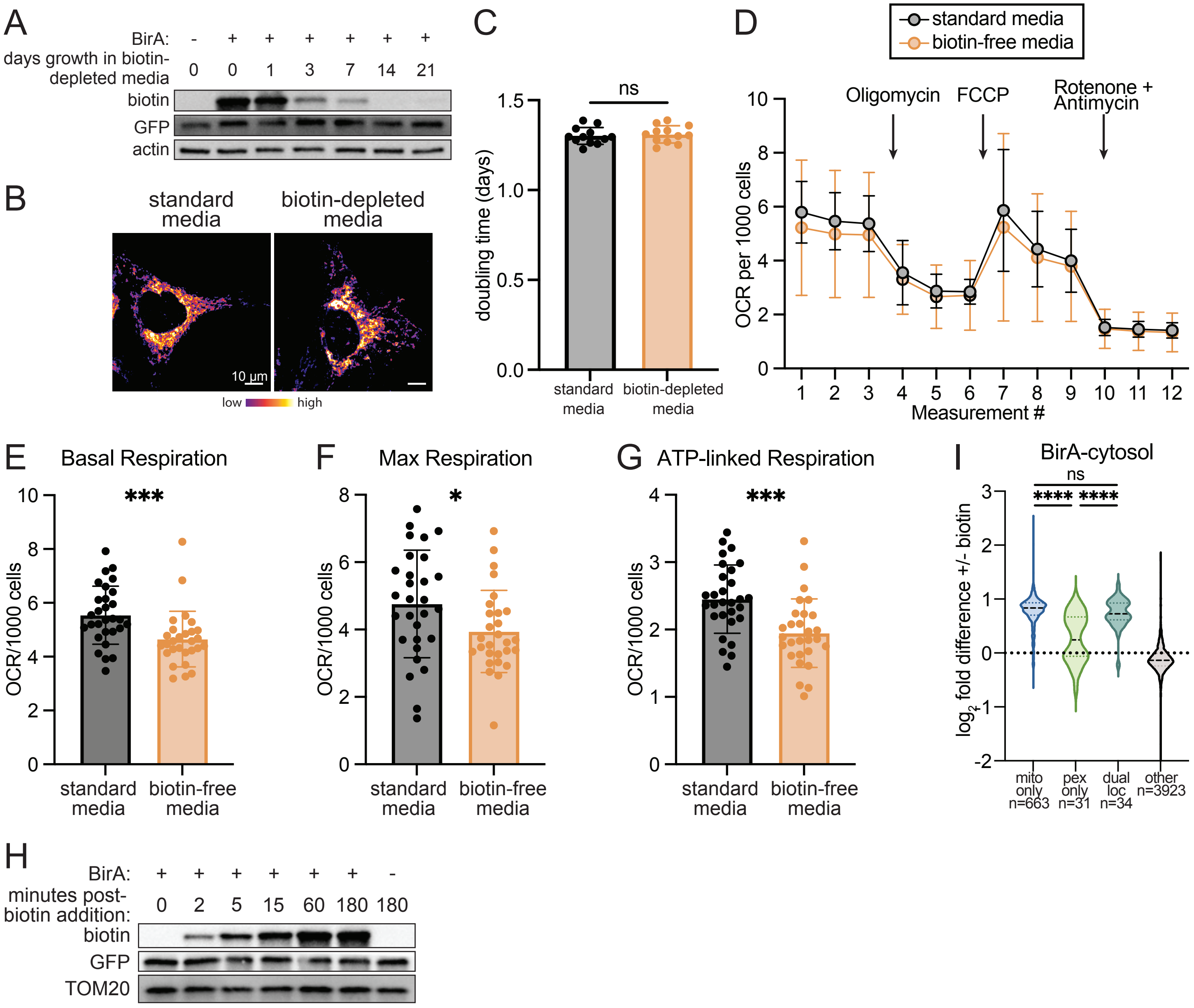


**Supplemental Figure 1: Biotin-dependent labeling and purification of mitochondria. (A)** Western blot analysis of whole cell-lysates harvested from U2OS cells expressing mitochondrial AviTag-GFP with or without cytosolic BirA-mCherry. Cells were grown in standard or biotin-depleted media for the indicated times. **(B)** Confocal microscopy of U2OS cells expressing AviTag-GFP grown in standard or biotin-depleted media for 3 weeks **(C)** Doubling time calculated for cells grown as in **B** and assessed over a 4-day period. **(D)** Oxygen consumption rate of cells grown as in **B** measured by Seahorse assay (n=24 across 3 biological replicates). **(E)** Basal respiration calculated using the data in **D**. **(F)** Maximal respiration calculated using the data in **D**. **(G)** ATP-linked respiration calculated using the data in **D**. **(H)** Western blot analysis of whole cell-lysate harvested from U2OS cells expressing mitochondrial AviTag-GFP with or without cytosolic BirA-mCherry. Cells were grown in biotin-depleted media for three weeks and treated with 50 µM biotin for the indicated times. **(I)** Violin plot showing the distribution of biotin-dependent enrichment values for proteins localized to the indicated organelles. “Dual loc” indicates proteins with annotated localization to both mitochondria and peroxisomes. Statistical comparisons for all panels are one-way ANOVA with correction for multiple comparisons. ns: $p>0.05$; *: $p<0.05$; ***: $p<0.001$; ****: $p<0.0001$.

### Supplemental Figure 2

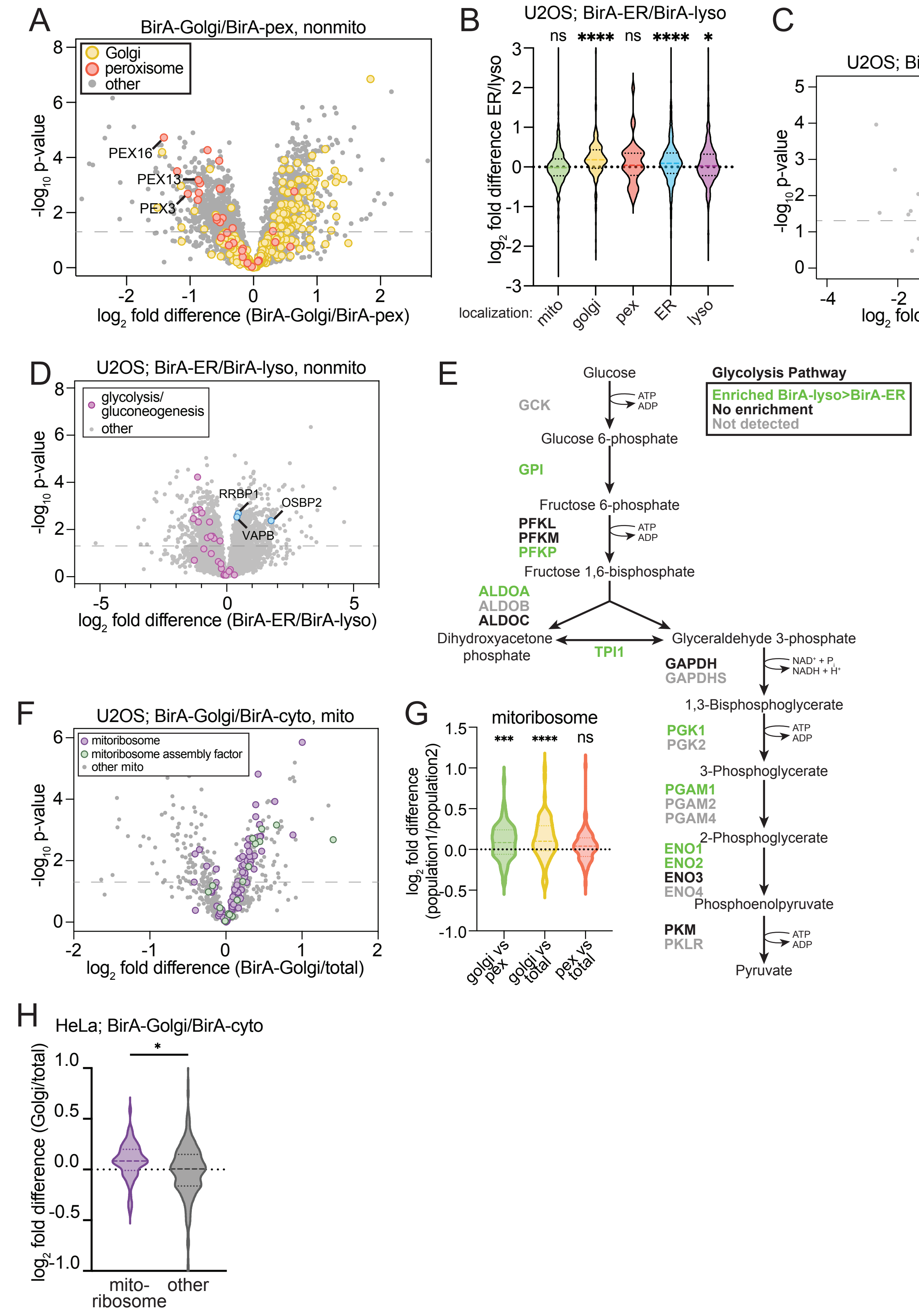

**Supplemental Figure 2: ORCA reveals proteomic differences between mitochondrial subpopulations. (A)** Volcano plot showing relative non-mitochondrial protein abundance in Golgi- and peroxisome-associated mitochondria, with the indicated protein categories enriched in peroxisome-associated mitochondria highlighted. P-values were calculated by student's t-test. **(B)** Violin plot comparing proteins localized to the indicated organelles for their enrichment in ER-proximal and lysosome-proximal mitochondrial preparations. Samples were compared to a hypothetical value of 0 by one-sample t-test. **(C)** Volcano plot showing relative mitochondrial protein abundance in ER- and lysosome-associated mitochondria, with select proteins highlighted. P-values were calculated by student's t-test. **(D)** As in **C** but for non-mitochondrial proteins. **(E)** Diagram of the glycolysis pathway adapted from Cheng et. al. (2013) (PMID: 22834840) with protein names annotated based on their enrichment and detection. **(F)** Volcano plot showing relative mitochondrial protein abundance in Golgi-associated mitochondria relative to total mitochondrial pool, with select proteins highlighted. P-values were calculated by student's t-test. **(G)** Violin plot comparing the distribution of enrichment for all mitoribosomal proteins in mitochondrial preparations from U2OS cells expressing BirA-cytosol, BirA-Golgi, and BirA-pex. Statistical analysis shows comparison to a hypothetical value of 0 by one-sample t-test. **(H)** Violin plot comparing the distribution of enrichment for all mitoribosomal proteins in mitochondrial preparations from HeLa cells expressing BirA-cytosol and BirA-Golgi. Samples were compared by one-way ANOVA. Mito: mitochondria; pex: peroxisome; lyso: lysosome; ns: $p>0.05$; *: $p<0.05$; ***: $p<0.001$; ****: $p<0.0001$.

### Supplemental Figure 3

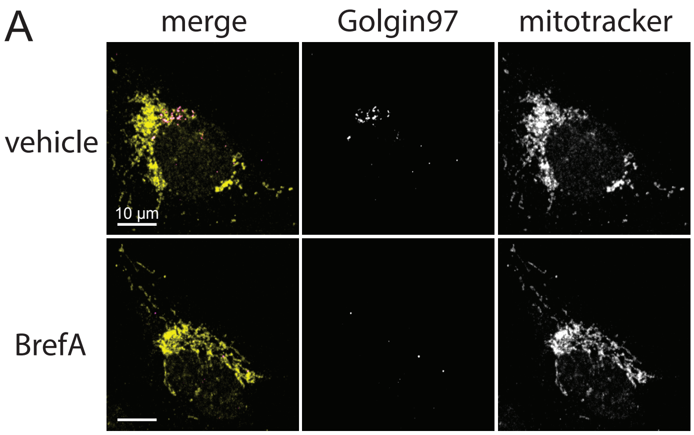


**Supplemental Figure 3: Mitochondrial translation is enriched at Golgi contact sites. (A)** Confocal microscopy of U2OS cells treated with 1 µg/mL Brefeldin A or an equivalent volume of DMSO for four hours and labeled with MitoTracker Deep Red FM and anti-Golgin-97.

### Supplemental Figure 4

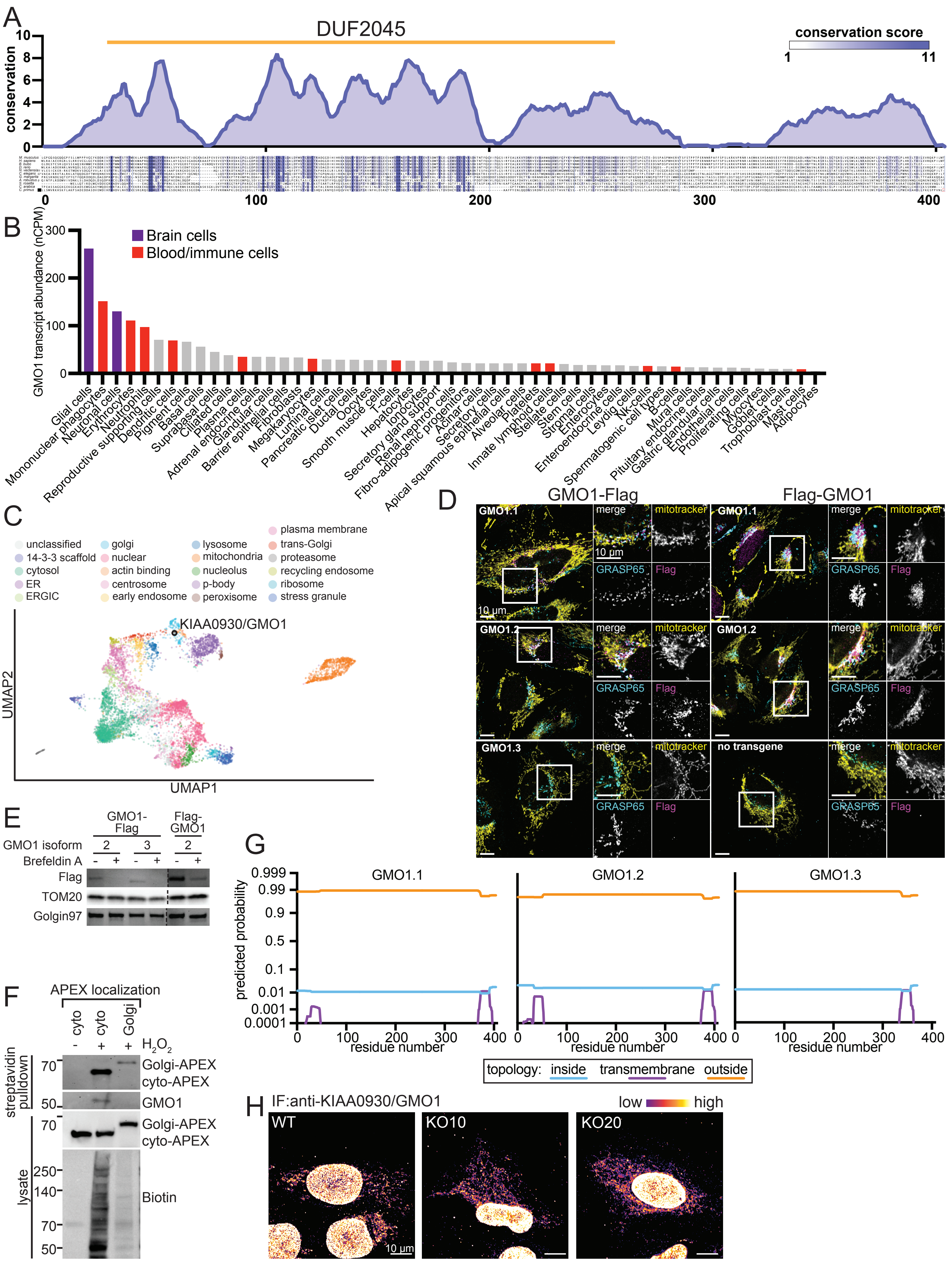

**Supplemental Figure 4: KIAA0930/GMO1 is a novel Golgi protein enriched at mitochondria contact sites. (A)** Conservation analysis of select GMO1 orthologs detailed in Table 1. Top: Conservation score represented as a 10-amino acid rolling average. Bottom: Sequence alignment generated using Clustal Omega and Jalview with each residue color coded according to its conservation score. **(B)** GMO1 transcript abundance across the indicated cell types. Data accessed from the Human Protein Atlas as detailed in Karlsson et. al. (50). **(C)** Two-dimensional UMAP generated by Hein et. al. (51) showing subcellular compartment clustering based on enrichment scores across a suite of organelle immunocapture experiments. Data accessed from organelles.czbiohub.org. **(D)** Confocal microscopy of U2OS cells expressing the indicated isoforms of GMO1 tagged at the C-terminus (left) or N-terminus (right) labeled with MitoTracker Deep Red FM and stained with Flag and Golgi protein GRASP65 antibodies. **(E)** Western blot analysis of whole cell lysates from U2OS cells expressing the indicated Flag-tagged GMO1 isoforms and treated with 1μg/mL Brefeldin A for 4 hours. **(F)** Western blot analysis of whole-cell lysates and streptavidin pulldowns from U2OS cells expressing the indicated APEX2 transgene with or without peroxide treatment. **(G)** Predicted probability of membrane topologies for each amino acid in the indicated GMO1 isoform. Data generated using Transmembrane Helix Prediction (TMHMM2.0) accessed at services.healthtech.dtu.dk/services/TMHMM-2.0/ (53). **(H)** Confocal microscopy of U2OS WT or GMO1 knockout cells labeled with an anti-GMO1 antibody. nCPM: normalized counts per million; DUF2045: Domain of Unknown Function 2045.

Supplemental Figure 5

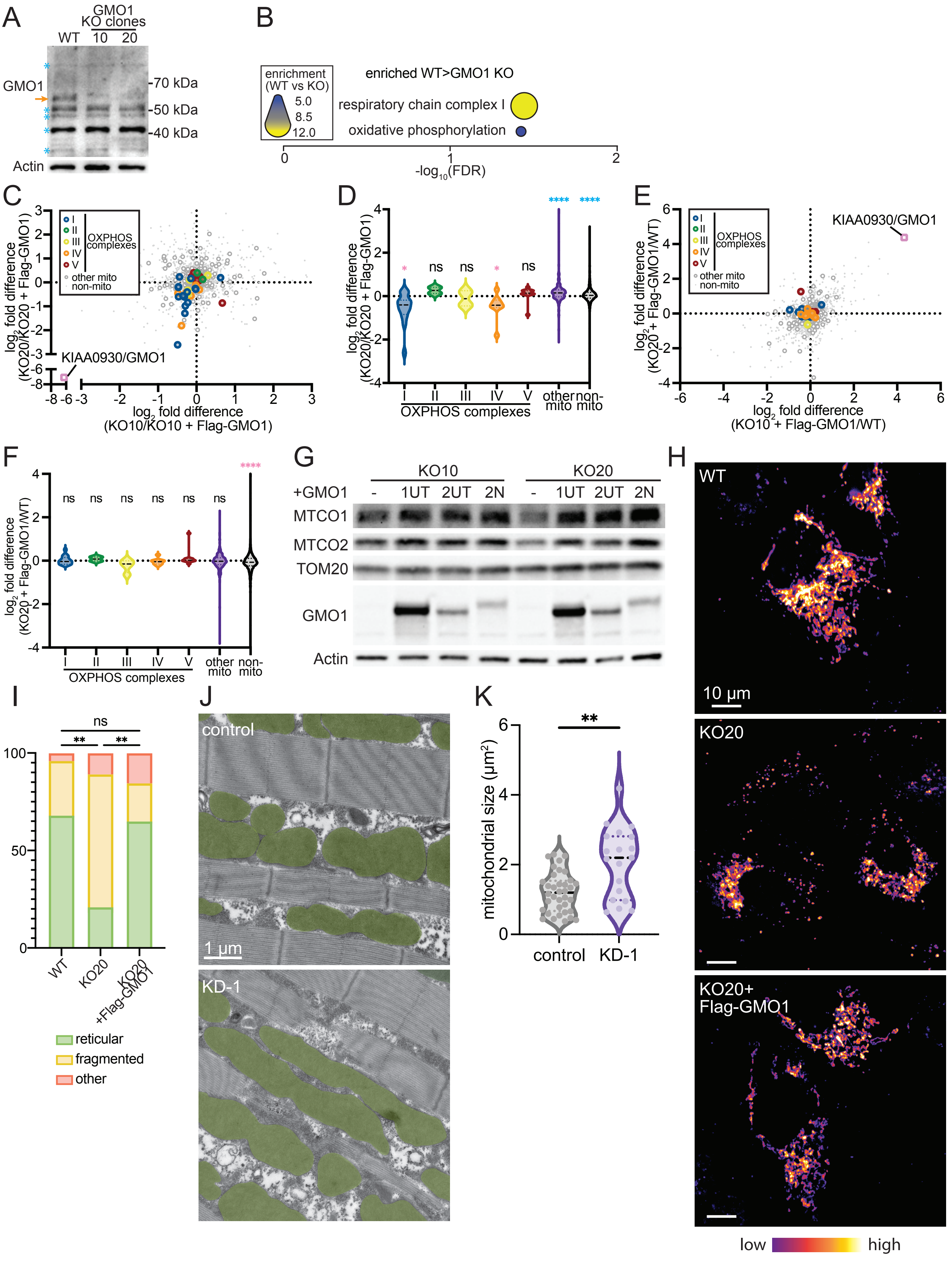

**Supplemental Figure 5: GMO1 is a conserved regulator of mitochondrial metabolism and morphology. (A)** Western blot analysis of U2OS WT or GMO1 knockout clones. Image is a wider crop of the image from **Figure 5A** illustrating abundant background bands. Orange arrow indicates the knockout-validated GMO1 band. Blue asterisk indicates background bands. **(B)** Gene set enrichment analysis showing fold-enrichment and FDR for all proteins enriched in WT U2OS cells compared to GMO1 knockout clones ($log_2$FC>0.5 for both WT vs KO10 and WT vs KO20). **(C)** Scatter plot showing the abundance for all proteins (n=4639) in U2OS GMO1 knockout clones relative to the same clones rescued by reintroduction of Flag-GMO1.2 with select protein categories highlighted. **(D)** Violin plot showing the distribution of fold-change in abundance for proteins in the indicated categories between GMO1 knockout clone 20 and the same clone rescued by reintroduction of Flag-GMO1.2. Each sample was compared to a hypothetical value of 0 by one-sample t-test, with pink and blue asterisks indicating significant downregulation or upregulation in GMO1 knockout cells, respectively. **(E)** As in **C** but comparing rescue cell lines to U2OS WT cells. **(F)** As in **D** but comparing GMO1 rescue clone 20 to U2OS WT cells. **(G)** Western blot analysis of whole-cell lysates from GMO1 knockout cells with and without re-expression of the indicated GMO1 isoform. **(H)** Confocal microscopy with cells of the indicated genotypes expressing mitochondrially targeted mCherry. **(I)** Categorization of mitochondria morphologies for cells imaged as in **H**. Conditions are compared via Chi-squared analysis. **(J)** Electron microscopy images of indirect flight muscle from flies expressing either CG9646-targeting RNAi or a non-targeting (w RNAi) control with mitochondria pseudocolored in green. **(K)** Quantification of mitochondrial area from images captured as in **J**. Conditions are compared by one-way ANOVA. ns: not significant (p>0.05); *: p<0.05; **: p<0.01; ****: p<0.0001. 1UT: GMO1.1 untagged; 2: 2UT: GMO1.2 untagged; 2N: Flag-GMO1.2.

**Supplemental Table 1: BirA targeting sequences**

| organelle | construct | source | citation |
|---|---|---|---|
| cytosol | BirA-mCherry-Flag | N/A | N/A |
| ER | ER-Flag-mCherry-BirA | rabbit cytochrome P450 2C1 | https://pubmed.ncbi.nlm.nih.gov/28441135/ |
| lysosome | LAMP1-Flag-mCherry-BirA | human LAMP1 | https://pubmed.ncbi.nlm.nih.gov/22053050/ |
| Golgi | BirA-mCherry-Flag-Golgi | human Golgin-97 | https://pubmed.ncbi.nlm.nih.gov/15136572/ |
| peroxisome | PEX3-Flag-mCherry-BirA | human PEX3 | https://pubmed.ncbi.nlm.nih.gov/10430017/ |

#### Supplemental Table 2: RNA FISH Probe sequences

**FISH IMAGER PROBE**
5´–ATGATGATGTATGATGATGTATGATGATGTTTTTTTTT–[AlexaFluor647]–3´

**Primary RNA FISH probes** (*listed 5´–>3´*)

**MT_RNR2**

TGTCTGGTAGTAAGGTGGAGTTTACATCATCATACATCATCATACATCATCAT
CCAGGTTTCAATTTCTATCGTTTACATCATCATACATCATCATACATCATCAT
TTGCGGTACTATATCTATTGTTTACATCATCATACATCATCATACATCATCAT
ATGCAGAAGGTATAGGGGTTTTTACATCATCATACATCATCATACATCATCAT
GCTCTCCTTGCAAAGTTATTTTTACATCATCATACATCATCATACATCATCAT
TTAGGTAGCTCGTCTGGTTTTTTACATCATCATACATCATCATACATCATCAT
CCCACTATTTTGCTACATAGTTTACATCATCATACATCATCATACATCATCAT
TGTCGCCTCTACCTATAAATTTTACATCATCATACATCATCATACATCATCAT
AAGGGGATTTAGAGGGTTCTTTTACATCATCATACATCATCATACATCATCAT
TGTTCCTCTTTGGACTAACATTTACATCATCATACATCATCATACATCATCAT
GTTTTTTCCTAGTGTCCAAATTTACATCATCATACATCATCATACATCATCAT
CTTAATTGGTGGCTGCTTTTTTTACATCATCATACATCATCATACATCATCAT
TAGTGGGTGTTGAGCTTGAATTTACATCATCATACATCATCATACATCATCAT
GTTCAGTTATATGTTTGGGATTTACATCATCATACATCATCATACATCATCAT
TAGATTGGTCCAATTGGGTGTTTACATCATCATACATCATCATACATCATCAT
CATTAGTTCTTCTATAGGGTTTTACATCATCATACATCATCATACATCATCAT
TGTTTTAATCTGACGCAGGCTTTACATCATCATACATCATCATACATCATCAT
TGGGCTGTTAATTGTCAGTTTTTACATCATCATACATCATCATACATCATCAT
ATGACTTGTTGGTTGATTGTTTTACATCATCATACATCATCATACATCATCAT
TGTGTTGGGTTGACAGTGAGTTTACATCATCATACATCATCATACATCATCAT
TGCCTCTAATACTGGTGATGTTTACATCATCATACATCATCATACATCATCAT
AAACATGTGTCACTGGGCAGTTTACATCATCATACATCATCATACATCATCAT
CAGGTTTGGTAGTTTAGGACTTTACATCATCATACATCATCATACATCATCAT
GCCCCAACCGAAATTTTTAATTTACATCATCATACATCATCATACATCATCAT
AAGTCTTAGCATGTACTGCTTTTACATCATCATACATCATCATACATCATCAT
TAGTAGTTCGCTTTGACTGGTTTACATCATCATACATCATCATACATCATCAT
CGTTGGTCAAGTTATTGGATTTTACATCATCATACATCATCATACATCATCAT
TTATCCCTAGGGTAACTTGTTTTACATCATCATACATCATCATACATCATCAT
ACTCTAGAATAGGATTGCGCTTTACATCATCATACATCATCATACATCATCAT
GTCGTAAACCCTATTGTTGATTTACATCATCATACATCATCATACATCATCAT
GATGTCCTGATCCAACATCGTTTACATCATCATACATCATCATACATCATCAT
CGAACCTTTAATAGCGGCTGTTTACATCATCATACATCATCATACATCATCAT
CGTAGGACTTTAATCGTTGATTTACATCATCATACATCATCATACATCATCAT
TACTCCGGTCTGAACTCAGATTTACATCATCATACATCATCATACATCATCAT
GTAGATAGAAACCGACCTGGTTTACATCATCATACATCATCATACATCATCAT
TGTGAAGTAGGCCTTATTTCTTTACATCATCATACATCATCATACATCATCAT
TATCATTTACGGGGGAAGGCTTTACATCATCATACATCATCATACATCATCAT
GTGGGTGTGGGTATAATACTTTTACATCATCATACATCATCATACATCATCAT

**Supplemental Table 3: List of antibodies used**

| primary antibodies | manufacturer | catalog # | application | dilution factor |
|---|---|---|---|---|
| biotin | Santa Cruz Biotechnology | sc-101339 | western | 1:1000 |
| GFP | Cell Signaling Technology | 2956 | western | 1:1000 |
| TOM20 | Cell Signaling Technology | 42406 | western | 1:1000 |
| OGDH | Abcam | ab137773 | western | 1:1000 |
| Calreticulin | Cell Signaling Technology | 12238 | western | 1:1000 |
| Golgin-97 | Cell Signaling Technology | 13192 | western, immunofluorescence | 1:1000 (W); 1:500 (IF) |
| LAMTOR1 | Cell Signaling Technology | 8975 | western, immunofluorescence | 1:1000 (W); 1:500 (IF) |
| Pex14 | Proteintech | 10594-1-AP | western, immunofluorescence | 1:1000 (W); 1:500 (IF) |
| Actin | Cell Signaling Technology | 3700 | western | 1:2000 |
| Pex19 | Abcam | ab137072 | immunofluorescence | 1:500 |
| Calnexin | Abclonal | A27819 | immunofluorescence | 1:500 |
| MRPL9 | Thermo Fisher | PA5-52581 | immunofluorescence | 1:500 |
| MT-CO1/COX1 | Abcam | ab14705 | western | 1:1000 |
| COX4 | Cell Signaling Technology | 4850 | western | 1:2000 |
| GRASP65 | Thermo Fisher | MA5-25148 | immunofluorescence | 1:500 |
| KIAA0930 | Thermo Fisher | PA5-44293 | western | 1:1000 |
| KIAA0930 | Sigma Aldrich | HPA038091 | immunofluorescence | 1:150 |
| OXPHOS | Proteintech | PK30006 | western | 1:1000 |
| Flag | Cell Signaling Technology | 14793 | western, immunofluorescence | 1:1000 (W); 1:500 (IF) |
| ATP5A | Abcam | ab14748 | immunofluorescence | 1:200 |

| secondary antibodies | manufacturer | catalog # | application | dilution factor |
|---|---|---|---|---|
| mouse-HRP | Cell Signaling Technology | 7076 | western | 1:5000 |
| rabbit-HRP | Cell Signaling Technology | 7074 | western | 1:5000 |
| mouse-AF488 | Thermo Fisher | A11011 | immunofluorescence | 1:2000 |
| rabbit-AF488 | Thermo Fisher | A32731 | immunofluorescence | 1:2000 |
| mouse-AF555 | Thermo Fisher | A28180 | immunofluorescence | 1:2000 |
| rabbit-AF555 | Thermo Fisher | A21428 | immunofluorescence | 1:2000 |
| rabbit-AF648 | Thermo Fisher | A21245 | immunofluorescence | 1:2000 |
